# Cylicins are a structural component of the sperm calyx being indispensable for male fertility in mice and human

**DOI:** 10.1101/2022.12.20.521208

**Authors:** Simon Schneider, Andjela Kovacevic, Michelle Mayer, Ann-Kristin Dicke, Lena Arévalo, Sophie A. Koser, Jan N. Hansen, Samuel Young, Christoph Brenker, Sabine Kliesch, Dagmar Wachten, Gregor Kirfel, Timo Strünker, Frank Tüttelmann, Hubert Schorle

## Abstract

Cylicins are testis-specific proteins, which are exclusively expressed during spermiogenesis. In mice and humans, two Cylicins, the gonosomal X-linked Cylicin 1 (*Cylc1/CYLC1*) and the autosomal Cylicin 2 (*Cylc2/CYLC2)* genes have been identified. Cylicins are cytoskeletal proteins with an overall positive charge due to lysine-rich repeats. While Cylicins have been localized in the acrosomal region of round spermatids, they resemble a major component of the calyx within the perinuclear theca at the posterior part of mature sperm nuclei. However, the role of Cylicins during spermiogenesis has not yet been investigated. Here, we applied CRISPR/Cas9-mediated gene-editing in zygotes to establish *Cylc1-* and *Cylc2*-deficient mouse lines as a model to study the function of these proteins. *Cylc1* deficiency resulted in male subfertility, whereas *Cylc2*^−/−^, *Cylc1*^−/y^ *Cylc2*^+/−^, and *Cylc1*^−/y^ *Cylc2*^−/−^ males were infertile. Phenotypical characterization revealed that loss of Cylicins prevents proper calyx assembly during spermiogenesis. This results in decreased epididymal sperm counts, impaired shedding of excess cytoplasm, and severe structural malformations, ultimately resulting in impaired sperm motility. Furthermore, exome sequencing identified an infertile man with a hemizygous variant in *CYLC1* and a heterozygous variant in *CYLC2*, displaying morphological abnormalities of the sperm including the absence of the acrosome. Thus, our study highlights the relevance and importance of Cylicins for spermiogenic remodeling and male fertility in human and mouse, and provides the basis for further studies on unraveling the complex molecular interactions between perinuclear theca proteins required during spermiogenesis.

## Introduction

The differentiation of round spermatids into sperm during spermiogenesis is a highly organized and spatiotemporally controlled process taking place in the seminiferous epithelium of the testis. Cellular and morphological remodeling involves DNA hypercondensation, establishment of a species-specific head morphology, removal of excess cytoplasm as well as formation of accessory structures like acrosome and flagellum. These structural changes depend on a highly efficient protein trafficking machinery and a unique sperm cytoskeleton ^1^. One essential cytoskeletal element is the perinuclear theca (PT), which surrounds the sperm nucleus, except for the caudal edge, at the implantation site of the flagellum. The PT is supposed to serve as a structural scaffold for the sperm nucleus and resembles a rigid cytosolic protein layer, which is resistant to non-ionic detergents and high salt buffer extractions ^2, 3^. The PT has been subdivided into a subacrosomal and postacrosomal part based on its localization, function, composition, and developmental origin. The subacrosomal part of the PT develops early during spermiogenesis, simultaneously with the formation of the acrosome. It is supposed to emerge from acrosomal vesicles, and presents as a thin cytosolic protein layer between the inner acrosomal membrane and the nuclear envelope ^4, 5^. The postacrosomal part of the PT, also known as the sperm calyx, originates from cytosolic proteins that are transported via the manchette and assembled during sperm head elongation ^4^. It is located in between the nuclear envelope and the sperm plasma membrane.

Proteomic analyses using murine and bovine PT extracts revealed 500-800 different proteins, highlighting its high molecular complexity ^6, 7^. Apart from cytoskeletal proteins, signaling molecules and several *de novo* synthesized core histones were identified. They are proposed to be essential for sperm-egg interaction and chromatin remodeling of the male pronucleus at early post-fertilization stages ^8, 9, 4, 10, 11^. Many structural proteins of the PT are testis-specific and uniquely expressed in the PT, including Calicin (*Ccin*) ^12, 2, 13^, Cylicin 1 (*Cylc1*) and Cylicin 2 (*Cylc2*) ^2, 14, 15^, actin-capping proteins CPβ3 and CPα3 ^16–18^ as well as actin-related proteins Arp-T1 and Arp-T2 ^19^. Despite being characterized on molecular level, their function remains largely elusive.

Both, *Ccin* and Cylicins are highly basic proteins with a predominant localization in the calyx of mature sperm ^14, 15, 12^. In most species, two Cylicin genes, *Cylc1* and *Cylc2*, have been identified (Figure 1-supplement 1). They are characterized by repetitive lysine-lysine-aspartic acid (KKD) and lysine-lysine-glutamic acid (KKE) peptide motifs, resulting in an isoelectric point (IEP) > pH 10 ^14, 15^. Repeating units of up to 41 amino acids in the central part of the molecules stand out by a predicted tendency to form individual short α-helices ^14^. Mammalian Cylicins exhibit similar protein and domain characteristics, but CYLC2 has a much shorter amino-terminal portion than CYLC1 (Figure 1 - supplement 1). While the *CYLC2/Cylc2* gene is encoded on autosomes, the human and murine *CYLC1/Cylc1* gene is encoded on the X-chromosome, resulting in hemizygosity in males. Further, in bovine, Cylc2 serves as actin-binding protein of the sperm perinuclear cytoskeleton ^20^. Cylicins seem to be cytoskeletal regulators and required for proper sperm head architecture.

**Figure 1:**
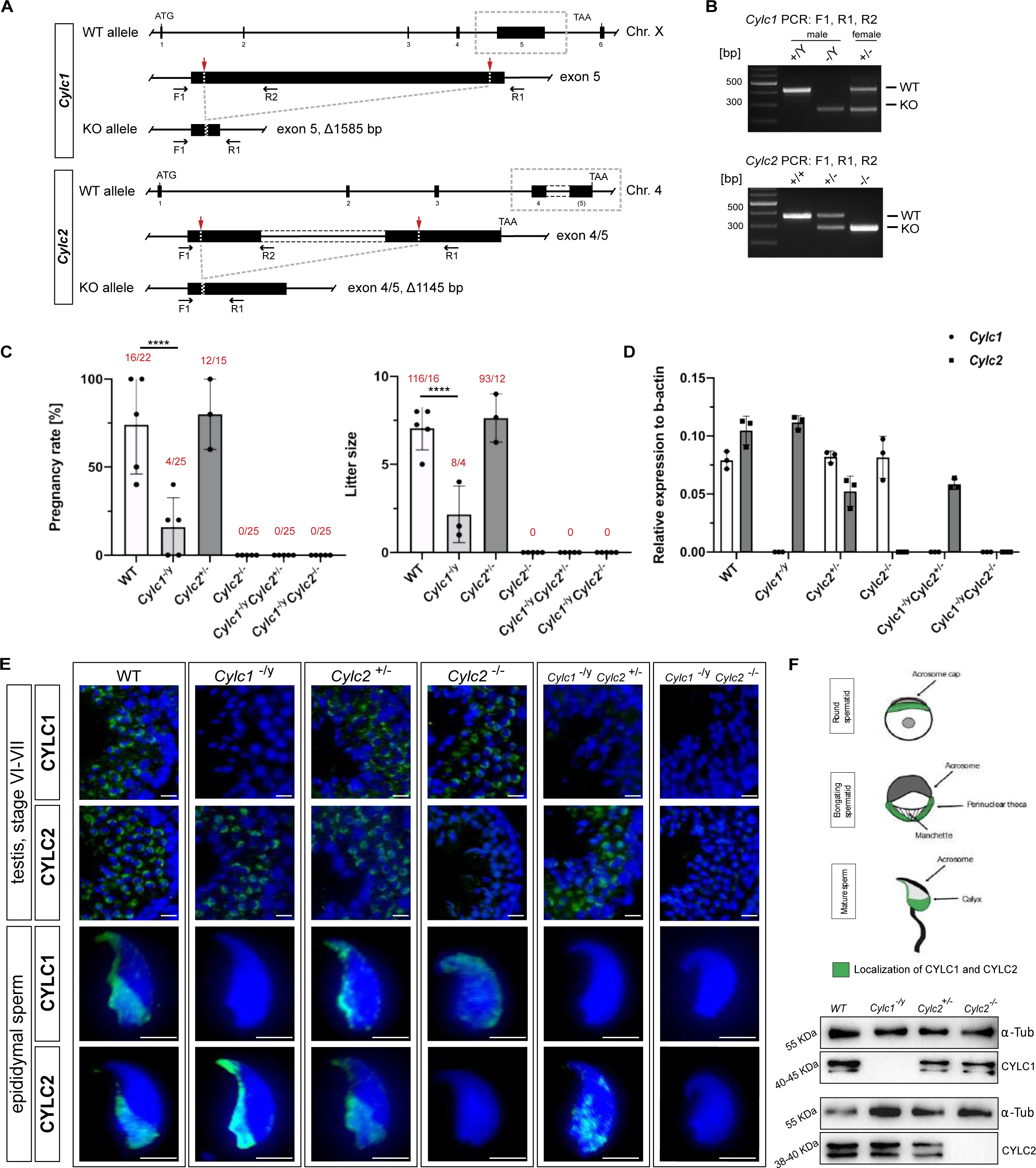
Loss of *Cylc1* or *Cylc2* results in impaired male fertility. (A) Schematic representation of the *Cylc1* and *Cylc2* gene structure and targeting strategy for CRISPR/Cas9-mediated generation of *Cylc1-* and *Cylc2-*deficient alleles. Targeting sites of guide RNAs are depicted by red arrows. Genotyping primer binding sites are depicted by black arrows. (B) Representative genotyping PCR of *Cylc1-* and *Cylc2-*deficient mice. N=3 (C) Fertility analysis of Cylicin-deficient mice visualized by mean litter size and pregnancy rate (%) in comparison to wildtype matings. Black dots represent mean values obtained for each male included in fertility testing. Columns represent mean values +/− standard deviation (SD). Total number of offspring per total number of pregnancies as well as total number of pregnancies per total number of plugs are depicted above each bar. (D) Expression of *Cylc1* and *Cylc2* in testicular tissue of WT, *Cylc1*^−/y^, *Cylc2*^+/−^, *Cylc2*^−/−^, *Cylc1*^−/y^ *Cylc2*^+/−^ and *Cylc1*^−/y^ *Cylc2*^−/−^ mice analyzed by qRT PCR. Biological replicate of 3 was used. (E) Immunofluorescent staining of testicular tissue and cauda epididymal sperm from WT, *Cylc1*^−/y^, *Cylc2*^+/−^, *Cylc2*^−/−^, *Cylc1*^−/y^ *Cylc2*^+/−^ and *Cylc1*^−/y^ *Cylc2*^−/−^ males against CYLC1 and CYLC2. Cell nuclei were counterstained with DAPI. Staining was performed on 3 animals from each genotype. Scale bar: 5 µm. (F) Schematic illustration of CYLC localization during spermiogenesis. CYLC localization (green) is shown for round and elongating spermatids as well as mature sperm. (G) Representative immunoblot against CYLC1 and CYLC2 on cytoskeletal protein fractions from WT, *Cylc1*^−/y^, *Cylc2*^+/−^ and *Cylc2*^−/−^ testes. α-Tubulin was used as load control.

In this study, we report the CRISPR/Cas9-mediated generation and characterization of *Cylc1-*, *Cylc2-,* and *Cylc1/2*-deficient mice, demonstrating that Cylicins are indispensable for male fertility and play a key function in the formation of the sperm calyx. We show that Cylicins are required for the maintenance of the PT integrity during spermiogenesis and in mature sperm. Their deficiency results in morphological defects of the sperm head, acrosome, and mid-piece. Our analyses revealed that *Cylicin* genes are evolutionary conserved in rodents and primates. Furthermore, we identified Cylicin variants in an infertile man, demonstrating the conserved role of Cylicins in regulating male infertility.

## Results

### Cylicins are indispensable for male fertility in mice

Cylicins were first discovered and characterized in the early nineties due to their subcellular localization a role in sperm head architecture was postulated ^2, 14, 15^,. To address the question of the role of Cylicins during spermiogenesis, we used CRISPR/Cas9-mediated gene-editing to generate *Cylc1*- and *Cylc2*-deficient mouse models. First, for *Cylc1*, a pair of sgRNAs targeting exon 5 was designed, and a mouse line with a frameshift inducing deletion of 1.585 kb, accounting for 85% of the *Cylc1* coding sequence, was established (Fig. 1 A). Next, two guide RNAs targeting exon 4/5 of the *Cylc2* gene were applied to establish a *Cylc2*-deficient mouse line with a 1.145 kb frameshift inducing deletion (Fig. 1 A). In both lines, the majority of predicted functional domains with repetitive, lysine-lysine-aspartic acid (KKD) and lysine-lysine-glutamic acid (KKE) peptide motifs were depleted. Deletion was confirmed by PCR-based genotyping (Fig. 1 B).

Fertility testing of *Cylc1*^−/y^ males revealed significantly reduced pregnancy rates (16 %) and mean litter size (2.2) (Fig. 1 C). *Cylc2*^−/−^ males were infertile, while *Cylc2*^+/−^ males showed no significant difference in fertility parameters compared to wildtype mice (Fig. 1 C). Additionally, established mouse lines were intercrossed to generate *Cylc1*^−/y^ *Cylc2*^+/−^ and *Cylc1*^−/y^ *Cylc2*^−/−^ males. Of note, *Cylc1*^−/y^ *Cylc2*^+/−^, and *Cylc1*^−/y^ *Cylc2*^−/−^ males were infertile (Fig. 1 C). This indicates that loss of *Cylc1* alone is partially tolerated, as suggested by subfertility of *Cylc1*^−/y^ males, whereas the additional loss of one *Cylc2* allele renders male mice infertile. Taken together, the results suggest, that two functional Cylicin alleles are required for male fertility.

qRT PCR confirmed the absence of *Cylc1* and/or *Cylc2* transcripts in Cylicin-deficient animals (Fig. 1 D). In *Cylc2*^+/−^ animals expression of *Cylc2* was reduced by 50 %. Neither loss of *Cylc1* nor *Cylc2* resulted in upregulation of *Cylc2* or *Cylc1*, respectively.

Next, due to the lack of commercial antisera, polyclonal antibodies against murine CYLC1 and CYLC2 were raised to visualize the localization of Cylicins during spermiogenesis. Specificity of antibodies was proven by immunohistochemical stainings (IHC), showing a specific signal in testis sections only, but not in any other organ tested (Figure 1 - supplement 2). Immunofluorescence staining of wildtype testicular tissue showed presence of both, CYLC1 and CYLC2 from the round spermatid stage onward (Fig. 1 E). The signal was first detectable in the subacrosomal region as a cap-like structure, lining the developing acrosome (Fig. 1 E-F, Figure 1 – supplement 3). As the spermatids elongate, CYLC1 and CYLC2 move across the PT towards the caudal part of the cell (Figure 1 – supplement 4). At later steps of spermiogenesis, the localization in the subacrosomal part of the PT faded, while it intensified in the postacrosomal calyx region (Fig. 1 E-F). Of note, the localization of CYLC1 and CYLC2 in the calyx of mature sperm has been reported in bovine and human. The generated antibodies did not stain testicular tissue and mature sperm of *Cylc1-* and *Cylc2*-deficient males, except for a very weak unspecific background staining in the lumen of seminiferous tubules and the residual bodies of testicular sperm (Fig. 1 E). Additionally, Western Blot analyses confirmed the absence of CYLC1 and CYLC2 in cytoskeletal protein fractions of the respective knockout (Fig. 1 G), thereby demonstrating i) specificity of the antibodies and ii) validating the gene knockout. Of note, the CYLC1 antibody detects a double band at 40-45 KDa. This is smaller than the predicted size of 74 KDa as, but both bands were absent in Cylc1^−/y^. Similarly, the CYLC2 Antibody detected a double band at 38-40 KDa instead of 66 KDa. Again, both bands were absent in *Cylc2*^−/−^. Next, Mass spectrometry analysis of cytoskeletal protein fraction of mature spermatozoa was performed detecting both proteins in WT but not in the respective knockout samples (Figure 1 – supplement 5; Figure 1 – supplement 6).

### Sperm morphology is severely altered in Cylicin-deficient mice

Next, spermiogenesis of Cylicin-deficient males was analyzed in detail. Gross testicular morphology as well as testicular weight were not significantly altered (Fig. 2 A, B). The testicular morphology appeared unaltered, with all stages of spermatogenesis being detectable in hematoxylin and eosin (HE) stained testicular sections (Figure 2 – supplement 1). However, a strong decline in cauda epididymal sperm counts was observed in all Cylicin-deficient males. For *Cylc1*^−/y^ and *Cylc2*^+/−^ males, a moderate reduction of 40-47% was determined, whereas *Cylc2*^−/−^ and *Cylc1*^−/y^ *Cylc2*^+/−^ displayed an approx. 65% reduction in epididymal sperm counts compared to wildtype mice (Fig. 2 C). In *Cylc1*^−/y^ *Cylc2*^−/−^ males, spermiogenesis was most impaired, as indicated by an 85% reduction of the sperm count (Fig. 2 C). Eosin-Nigrosin staining revealed that the viability of epididymal sperm from all genotypes was not severely affected (Fig. 2 D, Figure 2 – supplement 2). However, viability of *Cylc2*^−/−^ and *Cylc1*^−/y^ *Cylc2*^−/−^ sperm was significantly reduced by approx. 15% compared to wildtype sperm. (Fig. 2 D).

**Figure 2:**
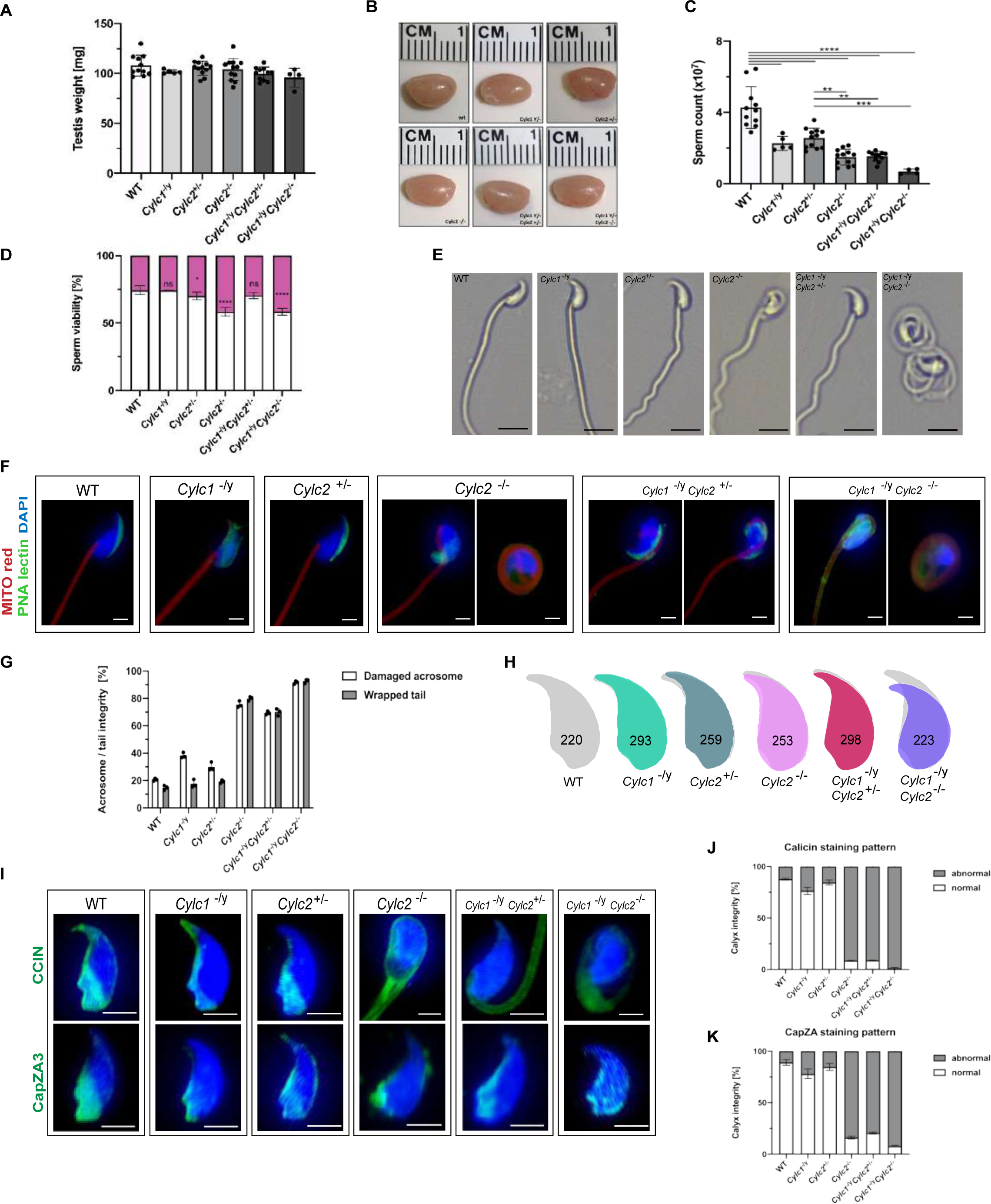
Sperm morphology is severely altered in Cylicin-deficient mice. (A) Testis weight [mg] and sperm count (x10^7^) of WT, *Cylc1*^−/y^, *Cylc2*^+/−^, *Cylc2*^−/−^, *Cylc1*^−/y^ *Cylc2*^+/−^ and *Cylc1*^−/y^ *Cylc2*^−/−^ males. Mean values +/− SD are shown; block dots represent data points for individual males. (B) Comparable photographs of the testes of WT, *Cylc1*^−/y^, *Cylc2*^+/−^, *Cylc2*^−/−^, *Cylc1*^−/y^ *Cylc2*^+/−^ and *Cylc1*^−/y^ *Cylc2*^−/−^ mice. (C) Epididymal sperm count (x10^7^) of WT, *Cylc1*^−/y^, *Cylc2*^+/−^, *Cylc2*^−/−^, *Cylc1*^−/y^ *Cylc2*^+/−^ and *Cylc1*^−/y^ *Cylc2*^−/−^ males. Mean values +/− SD are shown; block dots represent data points for individual males. (D) Viability of the epididymal sperm stained with Eosin Nigrosine. Percentage of Eosin negative (viable) and Eosin positive (inviable) sperm is shown. Data represented as mean +/− SD. Staining was performed on 3 animals from each genotype. (E) Bright field microscopy pictures of epididymal sperm from WT, Cylc1^−/y^, Cylc2^+/−^ and Cylc2^−/−^ mice. Scale bar: 10 μm. (F) Immunofluorescence staining of epididymal sperm acrosomes with PNA lectin (green) and tails with MITOred (red). Nuclei were counterstained with DAPI. Scale bar: 5 µm. (G) Quantification of abnormal sperm of WT, *Cylc1*^−/y^, *Cylc2*^+/−^, *Cylc2*^−/−^, *Cylc1*^−/y^ *Cylc2*^+/−^ and *Cylc1*^−/y^ *Cylc2*^−/−^ mice is shown. Acrosome aberrations and tail coiling were counted separately. Staining was performed on 3 animals from each genotype. (H) Nuclear morphology analysis of WT, *Cylc1*^−/y^, *Cylc2*^+/−^, *Cylc2*^−/−^, *Cylc1*^−/y^ *Cylc2*^+/−^ and *Cylc1*^−/y^ *Cylc2*^−/−^ sperm. Number of cells analyzed for each genotype is shown. (I) Representative pictures of immunofluorescent staining against PT proteins CCIN (upper panel) and CAPZa3 (lower panel) in WT, *Cylc1*^−/y^*, Cylc2*^+/−^, *Cylc2*^−/−^, *Cylc1*^−/y^ *Cylc2*^+/−^ and *Cylc1*^−/y^ *Cylc2*^−/−^ sperm. Nuclei were counterstained with DAPI. Staining was performed on 3 animals from each genotype. Scale bar: 5 µm. (J-K) Quantification of sperm with abnormal calyx integrity in WT, *Cylc1*^−/y^, *Cylc2*^+/−^, *Cylc2*^−/−^, *Cylc1*^−/y^ *Cylc2*^+/−^ and *Cylc1*^−/y^ *Cylc2*^−/−^ mice based on CCIN and CapZA staining patterns.

Next, we used brightfield microscopy to evaluate the effects of Cylicin deficiency on sperm morphology. Above all, coiling of the sperm tails and kinked sperm heads were observed in *Cylc2*^−/−^ and *Cylc1*^−/y^ *Cylc2*^−/−^ males (Fig. 2 E). To confirm this, we used PNA-FITC lectin immunofluorescence staining to analyze acrosome localization in mature sperm, MITOred to visualize mitochondria in the flagellum and DAPI to stain the nucleus (Fig. 2 F). Loss of *Cylc1* alone caused malformations of the acrosome in around 38% of mature sperm, while their flagellum appeared unaltered and properly connected to the head. *Cylc2*^+/−^ males showed normal sperm tail morphology with around 30% of acrosome malformations. *Cylc2*^−/−^ mature sperm cells displayed morphological alterations of head and mid-piece (Fig. 2 F-G). 76% of *Cylc2*^−/−^ sperm cells showed acrosome malformations, bending of the neck region, and/or coiling of the flagellum, occasionally resulting in its wrapping around the sperm head in 80% of sperm (Fig. 2 F). While 70% of *Cylc1*^−/Y^ *Cylc2*^+/−^ sperm showed these morphological alterations, around 92% of *Cylc1*^−/Y^ *Cylc2*^−/−^ sperm presented with coiled tail and abnormal acrosome (Fig. 2 F-G).

To analyze in detail the sperm head, we used Nuclear Morphology software on DAPI stained sperm samples. *Cylc1*^−/Y^ and *Cylc2*^+/−^ sperm showed no alterations of the nuclear shape when compared to wildtype (Fig. 2 H, Fig. S2 C, D). However, heads of *Cylc2*^−/−^ and *Cylc1*^−/Y^ *Cylc2*^−/−^ sperm appeared smaller, with shorter hooks and increased circularity of the nuclei as well as reduced elongation (Fig. 2 H, Fig. S2 D). Interestingly, *Cylc1*^−/Y^ *Cylc2*^+/−^ sperm heads appeared unaltered, suggesting that only *Cylc2* has a crucial role for sperm head shaping, and one functional *Cylc2* allele is sufficient to maintain the correct shape of the nucleus (Fig. 2 H, Figure 2 – supplement 3).

To study the effects of Cylicin deficiency on sperm calyx integrity and morphology, we analyzed the localization of other calyx-specific proteins, such as CCIN and CapZα3. In epididymal sperm, CCIN co-localize with both CYLC1 and CYLC2 in the calyx (Figure 2 – supplement 4). In *Cylc*1^−/y^ and *Cylc2*^+/−^ sperm, CCIN localization remained unchanged, being present in the calyx and in the ventral portion of PT as described previously ^7^. However, in 91% of *Cylc2*^−/−^ sperm, CCIN localized to the tail or in random parts of the sperm head (Fig. 2 I, J). In 91% of *Cylc1*^−/Y^ *Cylc2*^+/−^ and 98% of *Cylc1*^−/Y^ *Cylc2*^−/−^ sperm, the localization of CCIN was also significantly altered, with the signal mainly being present in the sperm tail. CapZα3 forms a heterodimer with CapZβ3, creating a functional complex that localizes in the calyx ^21^. Immunofluorescence stainings revealed that the localization of CapZα3 remained unchanged in *Cylc1*^−/Y^ and *Cylc2*^+/−^ mice compared to wildtype mice. In 84% of *Cylc2*^−/−^ sperm cells, CapZα3 localized in the caudal portion of the head but without regular calyx localization (Fig. 2 I, K). Interestingly, *Cylc1*^−/Y^ *Cylc2*^+/−^ mice showed less severe anomalies of the calyx and although CCIN was located almost exclusively in the tail, CapZα3 maintained the correct calyx localization in around 30% of sperm (Fig. 2 I). Finally, 92% of *Cylc1*^−/Y^ *Cylc2*^−/−^ spermatozoa showed CapZα3 localization across the sperm head without regular calyx shape. These results suggest that the loss of Cylicins impairs the formation of calyx and the correct localization of its components, which might contribute to morphological anomalies of the sperm described initially.

### *Cylc2*^−/−^ sperm cells have altered flagellar beat

Transmission electron microscopy (TEM) of epididymal sperm confirmed the severe structural defects observed by light microscopy (Fig. 3 A, Figure 3 – supplement 1): *Cylc2*^−/−^ sperm showed coiling of the tail and dislocation of the head-tail connecting piece from the basal plate, resulting in parallel positioning of head and tail (Fig. 3 A, white arrowheads). Furthermore, in *Cylc2*^−/−^ sperm, excess of cytoplasm was observed, located around the nucleus and coiled tail (Fig. 3 A). Anomalies of the head were observed at the level of the PT, while the nuclei appeared unaltered. In all *Cylc2*^−/−^ sperm cells, the posterior portion of PT – calyx was absent (Fig. 3 A, red arrowheads). Instead of surrounding the nucleus entirely, the PT in *Cylc2*^−/−^ sperm appeared interrupted, missing completely its caudal part. Further, we observed loosening of the peri-acrosomal region, which is not compact and adherent to the nucleus (Fig. 3 A, green arrowheads). On the contrary, *Cylc1*^−/y^ sperm cells appeared healthy, with intact PT, acrosome and calyx.

**Figure 3:**
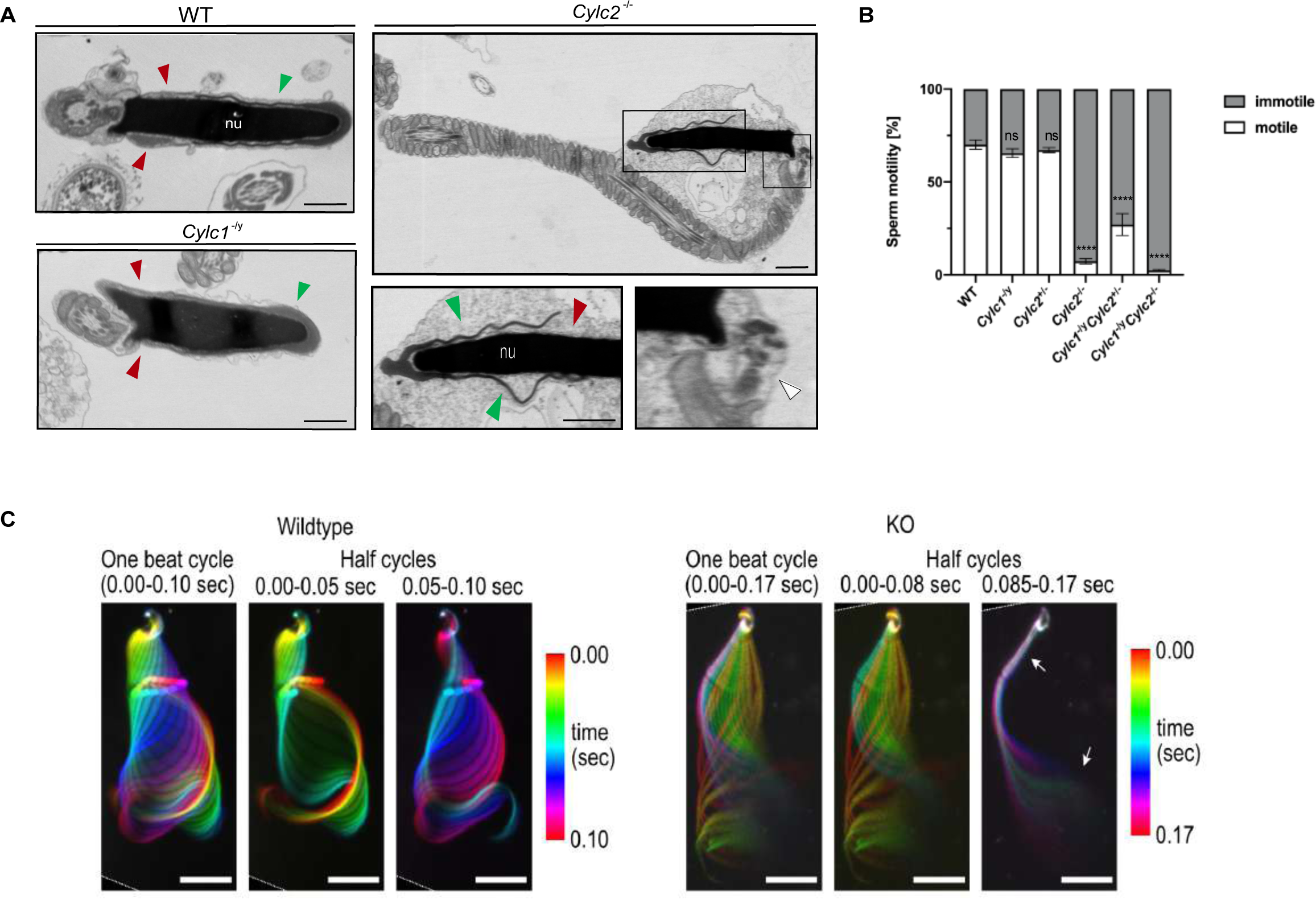
*Cylc2*^−/−^ sperm cells have altered flagellar beat. (A) TEM micrographs of WT, *Cylc1*^−/y^ and *Cylc2*^−/−^ epididymal sperm. Acrosome appears detached from the nucleus in *Cylc2*^−/−^ sperm (green arrowheads), while the calyx is missing entirely (red arrowheads). The head-tail connecting piece shifted from the basal plate is shown by white arrowheads causing the looping of the flagellum and formation of a cytoplasmatic sac. *Cylc1*^−/y^ sperm appears comparable to WT. Scale bar: 1 µm. (B) Motility of the epididymal sperm of WT, *Cylc1*^−/y^, *Cylc2*^+/−^, *Cylc2*^−/−^, *Cylc1*^−/y^ *Cylc2*^+/−^ and *Cylc1*^−/y^ *Cylc2*^−/−^ males activated in TYH medium. (C) Full and half-beat cycle plots of the flagellar beat are shown for WT and *Cylc2*^−/−^ spermatozoa. Half beat cycle shows the stiffness of the midpiece (upper arrow) and high oscillations (lower arrow) in *Cylc2*^−/−^ sperm in one direction of the beat.

While the motility of *Cylc1*^−/y^ and *Cylc2*^+/−^ sperm remained unchanged compared to wildtype sperm (around 60% motile cells), motility of *Cylc2*^−/−^ sperm was drastically reduced to only 7% motile sperm (Fig. 3 B) and the motility of *Cylc1*^−/y^ *Cylc2*^−/−^ sperm was reduced to 2% motile sperm. In addition, the few motile sperm cells were not progressive but were swimming in circular trajectories. Interestingly, in *Cylc1*^−/y^ *Cylc2*^+/−^ mice, sperm motility was reduced as well, but less drastically, with 27% of sperm cells being motile (Fig. 3 B).

The SpermQ software was used to analyze the flagellar beat of non-capacitated *Cylc2*^−/−^ sperm in detail ^22^. *Cylc2*^−/−^ sperm showed stiffness in the neck and a reduced amplitude of the initial flagellar beat, as well as reduced average curvature of the flagellum during a single beat (Figure 3 – supplement 2). Interestingly, we observed that the flagellar beat of *Cylc2*^−/−^ sperm cells was similar to wildtype cells, however, with interruptions during which midpiece and initial principal piece appeared stiff, whereas high frequency beating occurs at the flagellar tip (Fig. 3 C, Video 1, Video 2). These interruptions occurred only on the open-hook side and the duration of such interruptions varied from beat to beat. Of note, similar phenotypes have been observed for sperm with structural defects in the axoneme ^23^, however axoneme structure of Cylicin-deficient sperm appeared unaltered, presenting typical 9+2 microtubular composition in all genotypes (Figure 3 – supplement 1). Thus, we hypothesize that observed structural defects of the PT and head-tail connecting piece are restrictive for sperm motility and physiological beating patterns.

Taken together, observed anomalies of sperm heads, impaired sperm motility, and the decrease in epididymal sperm concentration show that Cylicin deficiency resembles a severe OAT (Oligo-Astheno-Teratozoospermia-syndrome) described in human.

### Cylicins are required for acrosome attachment to the nuclear envelope

To study the origin of observed structural sperm defects, spermiogenesis of Cylicin-deficient males was analyzed in detail. PNA lectin staining and Periodic Acid Schiff (PAS) staining of testicular tissue sections were performed to investigate acrosome biogenesis. During Golgi phase, the acrosome first starts to appear as an aggregation of proacrosomal vesicles into a single granule. This premature acrosomal structure was unaltered in all genotypes, with PNA signal appearing as a small dot on one pole of round spermatids (Fig. 4 A-B). During cap phase, acrosomes grow to cover the apical part of the nucleus. In wildtype and *Cylc2*^+/−^ mice, the forming acrosome appeared equally smooth and showed a regular cap structure on the perinuclear region of round spermatids. However, in some of the round spermatids from *Cylc2*^−/−^ and *Cylc1*^−/Y^ mice, gaps in the forming acrosome were observed, as well as an irregular shape of the cap. In *Cylc1*^−/Y^ *Cylc2*^+/−^ and *Cylc1*^−/Y^ *Cylc2*^−/−^ mice, most of the round spermatids were deformed or displayed irregularly localized caps (Fig. 4 A-B, Figure 4 – supplement 1, Figure 4 – supplement 2). At acrosome phase, many elongating spermatids of *Cylc1*^−/Y^, *Cylc2*^−/−^, *Cylc1*^−/Y^ *Cylc2*^+/−^ and *Cylc1*^−/Y^ *Cylc2*^−/−^ mice had irregular acrosome (Fig. 4 A-B, Figure 4 – supplement 1, Figure 4 – supplement 2). Detachment of the acrosome from the nuclear envelope was evident in testis samples of *Cylc2*^−/−^ and *Cylc1*^−/Y^ *Cylc2*^−/−^ male mice. These results suggest that Cylicins are required for the attachment of the developing acrosome to the nuclear envelope during spermiogenesis. Microtubules appeared longer on one side of the nucleus than on the other, displacing the acrosome to the side and creating a gap in the PT (Fig. 4 C). Whereas elongated spermatids at step 14-15 in wildtype sperm already disassembled their manchette and the PT appeared as a unique structure that compactly surrounds nucleus, in *Cylc2*^−/−^ spermatids, remaining microtubules failed to disassemble (Fig. 4 C, yellow arrowhead), and the acrosome detached from the nuclear envelope (green arrowhead). In addition, at step 16, the calyx was absent, and an excess of cytoplasm surrounded the nucleus and flagellum (Fig. 4 C, white arrowhead). Furthermore, many damaged and degrading cells were observed in testicular tissue TEM samples, having perforated nuclei and detached structures (Figure 4 – supplement 3). Interestingly, phagosomes with cellular remains were observed far away from the lumen and sometimes even at the basal membrane of the tubuli, suggesting that the cells that suffer most severe structural damage are being removed. This mechanism of removing malformed cells explains the reduction of epididymal sperm count in Cylicin-deficient genotypes.

**Figure 4:**
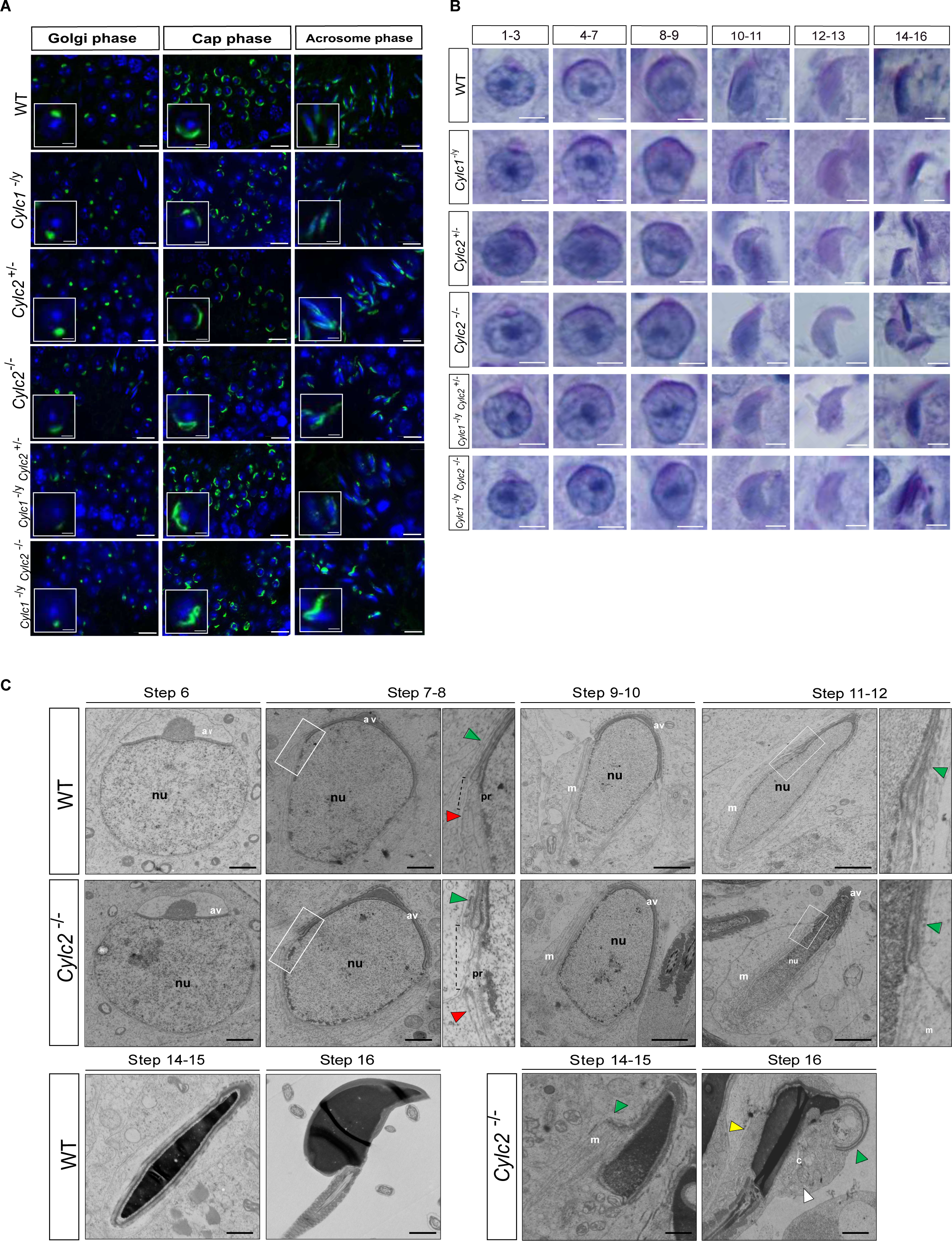
Cylicins are required for acrosome attachment to the nuclear envelope. (A) PNA-FITC lectin immunofluorescence staining of the acrosome in testicular tissue of WT, *Cylc1*^−/y^, *Cylc2*^+/−^, *Cylc2*^−/−^, *Cylc1*^−/y^ *Cylc2*^+/−^ and *Cylc1*^−/y^ *Cylc2*^−/−^ mice (green). Golgi phase of acrosome biogenesis at round spermatid stage (I-IV) is visible in the left panel. Middle panel shows cap phases on round spermatids (stage V-VIII). In the right panel acrosomal phase is shown (stage IX-XI). Nuclei were counterstained with DAPI. Staining was performed on 3 animals from each genotype. Scale bar: 10 µm. Insets show representative single spermatids at higher magnification (scale bar: 2 µm). (B) PAS staining of testicular sections from WT, *Cylc1*^−/y^, *Cylc2*^+/−^, *Cylc2*^−/−^, *Cylc1*^−/y^ *Cylc2*^+/−^ and *Cylc1*^−/y^ *Cylc2*^−/−^ mice. Representative spermatids at different steps of spermiogenesis are shown. Scale bar: 3 µm. (C) TEM micrographs of testicular tissues of WT and *Cylc2*^−/−^ mice. Single spermatids from step 6 to step 16 are shown. nu: nucleus; av: acrosomal vesicle; pr: perinuclear ring; m: manchette microtubules; cy: cytoplasm; green arrowheads: developing acrosome; red arrowheads: manchette; white arrowhead: cytoplasm; yellow arrowhead: remaining microtubules in mature sperm. Scale bar: 1 µm.

### Cylicin deficiency results in abnormal manchette elongation and disassembly

Next, we investigated the role of Cylicins during formation and development of the manchette – a sperm-specific, transient structure that represents a microtubular platform for protein transport, which showed several anomalies in TEM. Transport of the intracellular vesicles is crucial for the formation of the flagellum, acrosome assembly and removal of excess cytoplasm. The manchette is first detected at step 8 at the perinuclear ring of round spermatids, just prior to their elongation. During the next steps of spermiogenesis, as the spermatids elongate, manchette moves towards the neck region in a skirt-like structure and starts disassembling at step 13 when the elongation is complete ^24^. We used immunofluorescence staining of α-tubulin on squash testis samples containing spermatids at different stages of spermiogenesis to investigate whether the altered head shape, calyx structure, and tail-head connection anomalies originate from defects of the manchette structure. Spermatids starting from step 8 were observed individually for step-to-step comparison. In all genotypes, a cap-like shape of the manchette was observed in step 8 round spermatids, suggesting that the manchette assembles properly and starts elongating towards the neck region during step 9 (Figure 5 – supplement 1). In all genotypes, the typical skirt-like structure was observed at the caudal region of the spermatids at step 10 and 11, suggesting that the manchette assembles correctly even in Cylicin-deficient sperm (Fig. 5 A). In spermatids from *Cylc1*^−/Y^ and *Cylc2*^+/−^ mice, regular manchette development was observed in further steps of spermiogenesis (Fig. 5 A). However, starting from step 12, spermatids from *Cylc2*^−/−^, *Cylc1*^−/Y^ *Cylc2*^+/−^, and *Cylc1*^−/Y^ *Cylc2*^−/−^ mice showed abnormal manchette elongation, which became more prominent at step 13 (Fig. 5 A). Manchette length was measured starting from step 10 until step 13 spermatids and the mean was obtained, showing that the average manchette length was 76-80 nm in wildtype, *Cylc1*^−/Y^ and *Cylc2*^+/−^, while for *Cylc2*^−/−^ and *Cylc1*^−/Y^ *Cylc2*^−/−^ spermatids mean manchette length reached 100 nm (Fig. 5 B). *Cylc1*^−/Y^ *Cylc2*^+/−^ spermatids displayed an intermediate phenotype with a mean manchette length of 86 nm. Interestingly, some of *Cylc2*-deficient spermatids showed shifting of the manchette to the ventral side of the nucleus along with excessive elongation. At step 16, the manchette was normally disassembled in wildtype, *Cylc1*^−/Y^, and *Cylc2*^+/−^ spermatids (Fig. 5 A). However, *Cylc2* ^−/−^, *Cylc1*^−/Y^ *Cylc2*^+/−^, and *Cylc1*^−/Y^ *Cylc2*^−/−^ spermatids showed a persistent α-tubulin signal, indicating that disassembly of the manchette is delayed or incomplete (Fig. 5 A).

**Figure 5:**
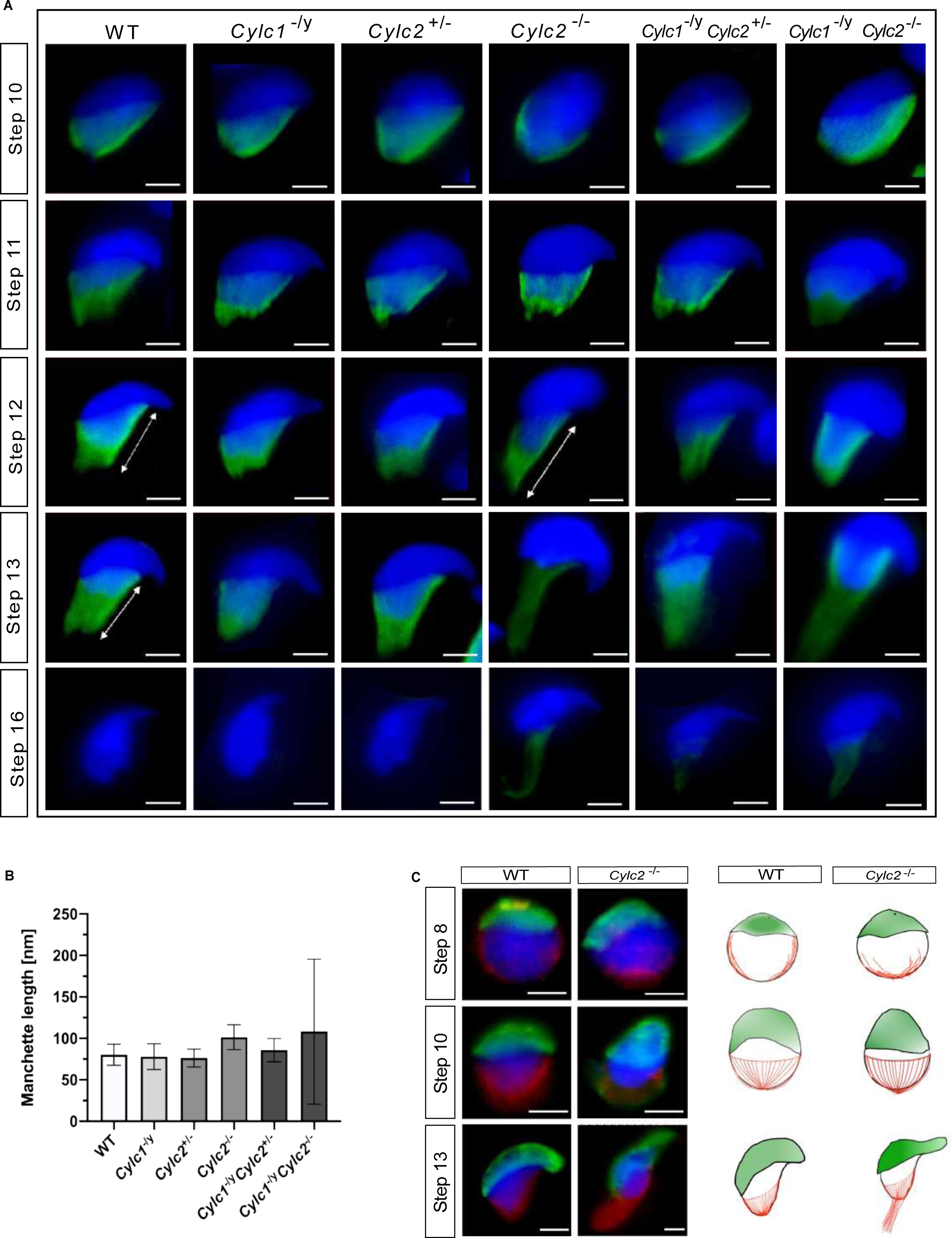
Cylc2 deficiency causes delay in manchette removal. (A) Immunofluorescence staining of α-Tubulin to visualize manchette structure in squash testis samples of WT, *Cylc1*^−/y^, *Cylc2*^+/−^, *Cylc2*^−/−^, *Cylc1*^−/y^ *Cylc2*^+/−^ and *Cylc1*^−/y^ *Cylc2*^−/−^ mice. Spermatids in different steps of spermiogenesis were shown, for step-to-step comparison. Scale bar: 5 µm. (B) Quantification of manchette length in WT, *Cylc1*^−/y^, *Cylc2*^+/−^, *Cylc2*^−/−^, *Cylc1*^−/y^ *Cylc2*^+/−^ and *Cylc1*^−/y^ *Cylc2*^−/−^ α-Tubulin stained spermatids at step 10-13. (C) Co-staining of the manchette with HOOK1 (red) and acrosome with PNA-lectin (green) is shown in round, elongating and elongated spermatids of WT (upper panel) and *Cylc2*^−/−^ mice (lower panel). Scale bar: 3 µm. Schematic representation shows acrosomal structure (green) and manchette filaments (red).

Other than α-tubulin, we also used HOOK1 as manchette marker. HOOK1 is a member of a family of coiled-coil proteins, which bind to microtubules and organelles and regulate microtubule trafficking during endocytosis and spermiogenesis. Co-staining of the spermatids with antibodies against PNA lectin (green) and HOOK1 (red) revealed that abnormal manchette elongation and acrosome anomalies simultaneously occurred in elongating spermatids of *Cylc2*^−/−^ male mice (Fig. 5 C). Schematic representation shows acrosome biogenesis and manchette development in wildtype and *Cylc2*^−/−^ spermatids (Fig. 5 C). While round spermatids of *Cylc2*^−/−^ mice elongated as those of the wildtype sperm, the manchette elongated abnormally and the acrosome became loosened (Fig. 4 C, Fig. 5 C).

### *Cylc2* coding sequence is slightly more conserved among species than *Cylc*1

To address why *Cylc2* deficiency causes more severe phenotypic alterations than *Cylc1* deficiency in mice, we performed evolutionary analysis of both genes. Analysis of the selective constrains on *Cylc1* and *Cylc2* across rodents and primates revealed that both genes’ coding sequences are conserved in general, although conservation is weaker in *Cylc1* trending towards a more relaxed constraint (Fig. 6). A model allowing for separate calculation of the evolutionary rate for primates and rodents, did not detect a significant difference between the clades, neither for *Cylc1* nor for *Cylc2*, indicating that the sequences are equally conserved in both clades.

**Figure 6:**
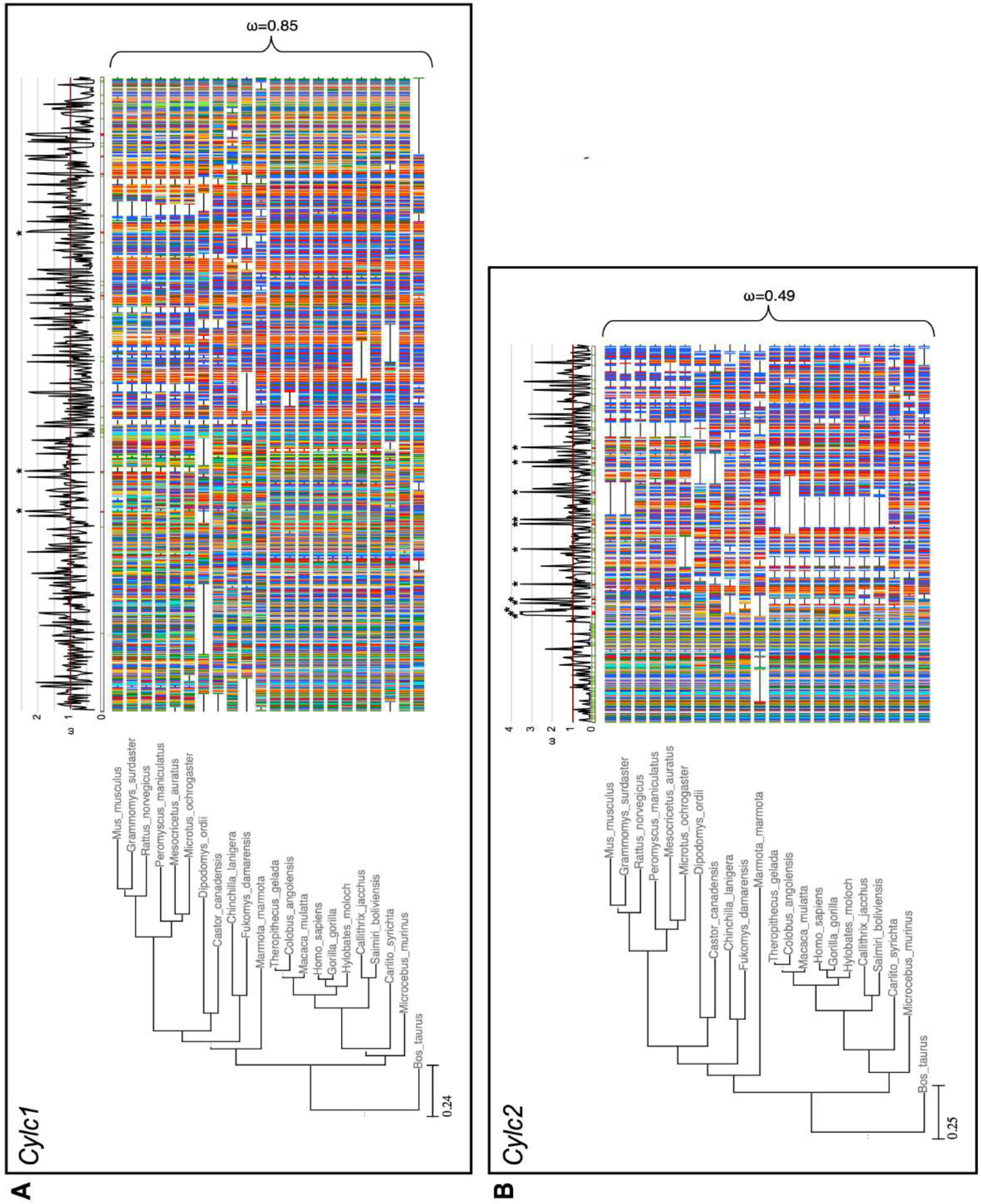
Species phylogeny with branch length representing number of nucleotide substitutions per codon with schematic representation of (A) CYLC1 and (B) CYLC2 amino acid alignment used in the PAML CodeML analysis. Alignments were stripped of columns with gaps in more than 80% of species. Evolutionary rate (ω) obtained by CodeML models M0 is shown for the whole alignment. The graph on top shows the evolutionary rate (ω) per codon sites across the whole tree (CodeML model M2a). Significantly positively selected sites are marked by asterisks. Sites with a probability of higher than 0.95 to belonging to the conserved or positively selected site class are marked in green and red respectively below the graph.

To analyze the selective pressure across the coding sequence in more detail, we calculated the evolutionary rates for each codon site across the whole tree. According to the analysis, 34% of codon sites were conserved, 51% under relaxed selective constraint, and 15% positively selected. For *Cylc2*, 47% of codon sites were conserved, 44% under neutral/relaxed constraint, and 9% positively selected. Interestingly, codon sites encoding lysine residues, which are proposed to be of functional importance for Cylicins, are mostly conserved. For *Cylc1*, 17% of lysine residues are significantly conserved and 35% of significantly conserved codons encode for lysine. For *Cylc2*, this pattern is even more pronounced with 27.9% of lysine codons being significantly conserved and 24.3% of all conserved sites encoding for lysine (Fig. 6).

### Cylicins are required for normal sperm morphology in human

As loss of two Cylicin alleles causes fertility defects in mice, we next addressed whether infertile men also display variants in *CYLC1/CYLC2*.

Exome sequencing within the MERGE (**M**al**e R**eproductive **G**enomics study) cohort identified one patient (M2270) carrying rare (MAF < 0.01, gnomAD) missense variants in both *CYLC1* and *CYLC2*. The man of German origin presented at age of 40 years for couple infertility because unsuccessfully trying to conceive for 6 years. The couple underwent one ICSI procedure which resulted in 17 fertilized oocytes out of 18 retrieved. Three cryo-single embryo transfers were performed in spontaneous cycles, but no pregnancy was achieved.

Patient M2270 carries the hemizygous variant c.1720G>C in *CYLC1* that leads to an amino acid exchange from glutamic acid to glutamine (p.(Glu574Gln)), is predicted to be deleterious or possibly damaging by *in silico* tools (SIFT ^25^ and PolyPhen ^26^, respectively), and has a CADD score of 11.91. It is located in exon four out of five and affects a region that is predicted to be intolerant to such substitutions (Figure 7 – supplement 1, metadome). It occurs only twice in the gnomAD database (v2.1.1) comprising 141,456 individuals (67,961 XY), once identified in a hemizygous male and once in a female carrier and is absent from our database.

**Figure 7:**
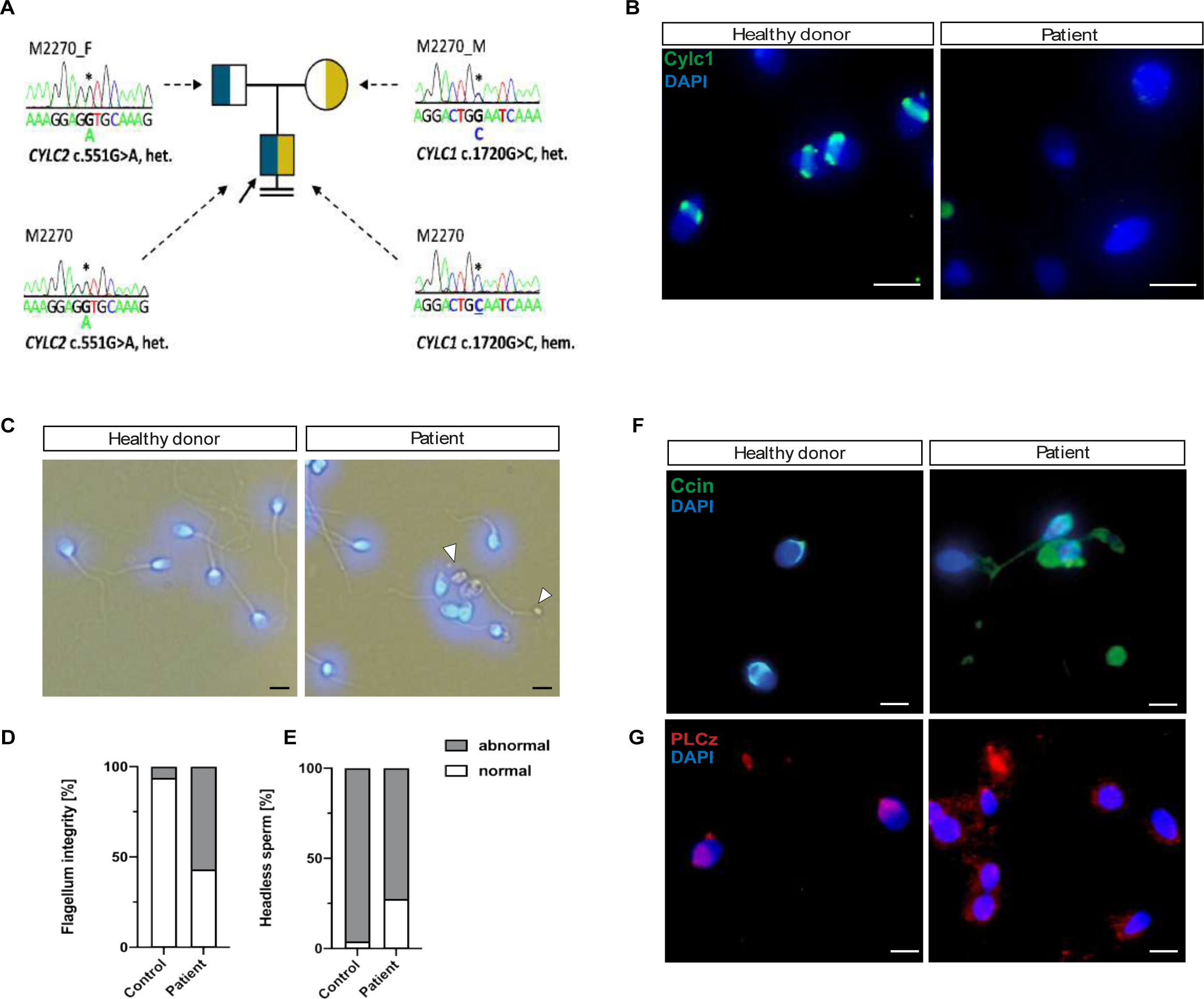
Cylicins are required for human male fertility. (A) Pedigree of patient M2270. His father (M2270_F) is carrier of the heterozygous *CYLC2* variant c.551G>A and his mother (M2270_M) carries the X-linked *CYLC1* variant c.1720G>C in a heterozygous state. Asterisks (*) indicate the location of the variants in *CYLC1* and *CYLC2* within the electropherograms. (B) Immunofluorescence staining of CYLC1 in spermatozoa from healthy donor and patient M2270. In donor’s sperm cells CYLC1 localizes in the calyx, while patient’s sperm cells are completely missing the signal. Scale bar: 5 µm. (C) Bright field images of the spermatozoa from healthy donor and M2270 show visible head and tail anomalies, coiling of the flagellum as well as headless spermatozoa, which carry cytoplasmatic residues without nuclei (white arrowhead). Heads were counterstained with DAPI. Scale bar: 5 µm. (D-E) Quantification of flagellum integrity (D) and headless sperm (E) in the semen of patient M2270 and a healthy donor. (F-G) Immunofluorescence staining of CCIN (F) and PLCz (G) in sperm cells of patient M2270 and a healthy donor. Nuclei were counterstained with DAPI. Scale bar: 3 µm.

M2270 further carries the heterozygous variant c.551G>A in *CYLC2* that is predicted to be tolerated (SIFT) or benign (PolyPhen) in accordance with a low CADD score of 0.008. It is located in exon five out of eight and affects a region in which variants are likely to be tolerated (Figure 7 – supplement 1, metadome). However, it is a rare variant occurring with an allele frequency of 0.0035 in the general population, according to the gnomAD database. Importantly, only three XX individuals are reported to be homozygous for the variant within *CYLC2*.

Segregation analyses revealed maternal inheritance of the X-linked *CYLC1* variant c.1720G>C p. (Glu574Gln), whereas the father carries the heterozygous *CYLC2* variant c.551G>A p.(Gly184Asp) (Fig. 7 A). According to ACMG-AMP criteria ^27^ both variants are classified as variants of uncertain significance. No other potentially pathogenic variants in genes associated with sperm morphological defects were identified by exploring the exome data of M2270.

Semen analysis performed following WHO guidelines ^28^ is shown in Table 1. The sperm concentration in the semen was slightly reduced, while significant reduction of motile spermatozoa (12.5%) was observed. Interestingly, only 2% of the sperm cells appeared morphologically normal, while 98% of sperm showed head defects (Table 1). Immunofluorescence staining of CYLC1 revealed that while in healthy donor’s sperm, CYLC1 localizes in the calyx, in M2270, CYLC1 labeling was absent (Fig. 7 B). Bright field microscopy demonstrated that M2270’s sperm flagella coiled in a similar manner compared to flagella of sperm from Cylicin-deficient mice. Quantification revealed 57% of M2270 sperm displaying abnormal flagella compared to 6% in the healthy donor (Fig. 7 D). Interestingly, DAPI staining revealed that 27% of M2270 flagella carry cytoplasmatic bodies without nuclei and could be defined as headless spermatozoa (Fig. 7 C, white arrowhead; Fig. 7 E). CCIN staining demonstrated that while spermatozoa of a healthy donor showed a typical, funnel-like calyx structure in the posterior region of PT, spermatozoa from M2270 had CCIN localized in irregular manner throughout head and tail, suggesting that Cylicins have a role in maintenance of the calyx structure and composition in both mice and human spermatozoa (Fig. 7 F). Testis-specific phospholipase C zeta 1 (PLCζ1) is localized in the postacrosomal region of PT in mammalian sperm ^29^ and has a role in generating calcium (Ca²⁺) oscillations that are necessary for oocyte activation ^30^. Staining of healthy donor’s spermatozoa showed a previously described localization of PLCζ1 in the calyx, while sperm from M2270 patient presents signal irregularly through the PT surrounding sperm heads (Fig. 7 G). These results suggest that Cylicin deficiency can cause severe disruption of PT in human sperm as well, causing male infertility.

**Table 1:** Semen analysis of the patient M2770 carrying variants in the *CYLC1* and *CYLC2* genes.

|  | First visit | Second visit | WHO reference rang |
| --- | --- | --- | --- |
| **Abstinence time [d]** | 4.0 | 5.0 |  |
| **Volume [ml]** | 4.2 | 5.8 | > 1.4 |
| **Concentration [Mill./ml]** | 10.5 | 16.3 | > 16 |
| **Total sperm count [Mill.]** | 44.1 | 94.5 | > 39 |
| **Vitality [%]** | 53 | 27 | > 54 |
| **Motility** |  |  |  |
| **a [%]** | 7 | 9 | a + b >30 |
| **b [%]** | 5 | 4 |  |
| **c [%]** | 19 | 8 |  |
| **d [%]** | 69 | 79 |  |
| **Morphology** |  |  |  |
| **Normal [%]** | 2 | 2 | > 4 |
| **Head defects [%]** | 99 | 99 |  |
| **Midpiece defects [%]** | 63 | 59 |  |
| **Flagella defects [%]** | 18 | 47 |  |

## Discussion

Spermiogenesis is a highly organized process that is dependent on a unique cytoskeletal organization of the sperm cells. The PT has a role of structural scaffold that surrounds sperm nucleus, and its protein composition is crucial for correct sperm development. In this study, we used CRISPR/Cas9 gene editing to establish *Cylc1-*, *Cylc2-* and *Cylc1/2*-deficient mouse lines to analyze the role of Cylicins. We demonstrated that the loss of Cylicins impairs male fertility in mice by severely disturbing sperm head architecture. The significance of our findings is supported by the identification of *CYLC1/2* variants in an infertile patient who presents similar structural anomalies in sperm cells.

Detailed analysis of sperm development and morphology demonstrated that infertility of Cylicin-deficient male mice is caused by anomalies of different sperm structures. The prominent loss of PT integrity and calyx structure in the mature sperm lead to acrosome loosening and detachment from the nuclear envelope. Furthermore, the shifting of the basal plate caused damage in the head-tail connecting piece resulting in coiling of the flagellum and impaired swimming. Murine Cylicins are first detected in the periacrosomal region of round spermatids and move to the postacrosomal region of PT during spermatid elongation to finally localize to the calyx of mature sperm. This dynamic localization pattern of Cylicins has been described in human, boar and bovine sperm as well ^2, 14, 15^.

Interestingly, other PT enriched proteins CCIN ^12, 31, 7^ and CPβ3/CPα3 move in the similar manner across PT during spermiogenesis and are present in the calyx of mature sperm of various mammalian species (Figure 2 – supplement 4). This co-localization suggests the potential interaction between calyx proteins. Furthermore, CCIN ^2, 12, 13^ and CPβ3-CPα3 complex ^16–18^ are described as actin-binding proteins and porcine Cylc2 has been shown to have a high affinity for F-actin as well ^20^. The potential roles of F-actin during spermiogenesis in mammals involve biogenesis of the acrosome ^32^ and its correct attachment to the outer nuclear membrane of the spermatids ^33^ as well as removal of excess cytoplasm ^34^.

The loss of Cylicins caused acrosome detachment from NE starting from cap phase of the acrosome biogenesis. Interestingly, loss of CCIN results in similar loosening of the acroplaxome from the outer nuclear membrane ^7^. CCIN is shown to be necessary for the IAM-PT-NE complex by establishing bidirectional connections with other PT proteins. Zhang et al. found CYLC1 to be among proteins enriched in PT fraction ^7^. Based on their speculation that CCIN is the main organizer of the PT, we hypothesize that both CCIN and Cylicins might interact, either directly or in a complex with other proteins, in order to provide the ‘molecular glue’ necessary for the acrosome anchoring. Furthermore, Cylicin deficiency resulted in cytoplasmatic retention and bending of the midpiece, similarly to the CPα3 mutant phenotype repro32/repro32 ^35^. The CAPZ complex has been shown to regulate the actin dynamics by preventing addition or loss of G-actin subunits to the ends of F-actin filaments ^36^. Since we demonstrated that the loss of Cylicins resulted in mislocalization of both CCIN and CPα3, we speculate that Cylicins have a crucial role in maintenance of the integrity of PT structure and thus are required for proper function of PT proteins.

Other than morphological defects of the mature sperm PT, during spermiogenesis Cylicin deficiency results in excessive manchette elongation, its delayed disassembly as well as formation of abnormal gaps in the PT at the level of perinuclear ring. The intra-manchette transport (IMT) of proteins from the apical pole of the head to the base of the developing tail occurs during spermiogenesis ^37^. The malfunctions of the IMT can cause delay in manchette clearance and morphological defects of mature sperm. Mouse models deficient for IMT proteins such as HOOK1 ^38^ CEP131 ^39^ and IFT88 ^40^ show abnormal manchette elongation and delay in its clearance resulting in aberrant nuclear shape of the sperm. Furthermore, the loss of HOOK1 results in head-tail connection and mid-piece anomalies such as flipping of the head, basal plate defects and coiling of the tail similar to *Cylc2* deficiency ^41^. The localization of manchette on the caudal portion of spermatids coincides with the localization of the calyx in the mature sperm, so these results indicate that the manchette might be maintained longer to compensate for the missing formation of the calyx structure. Furthermore, we observed wide gaps in the perinuclear ring during manchette elongation, suggesting that Cylicins might have a role in maintenance of the contact between caudal and apical PT region and its integrity.

Our evolutionary analysis of *Cylc1* and *Cylc2* genes across rodents and primates revealed that both coding sequences are under purifying selection. Overall, the results reveal that *Cylc1* is under slightly less conserved constraint than *Cylc2* leading to the hypothesis that loss of function in *Cylc1* might be less severe and could be compensated for by *Cylc2* due to partial redundancy. This hypothesis is supported by our finding that *Cylc1* deficiency causes subfertility in male mice, while the loss of both *Cylc2* alleles results in male infertility. Furthermore, *in Cylc2*^+/−^ male mice fertility was preserved, while *Cylc1*^−/Y^ *Cylc2*^+/−^ males were unable to sire offspring leading to the conclusion that the loss of one *Cylc2* allele could be compensated by *Cylc1*, however at least two functional Cylicin alleles are required for male fertility in mice. Interestingly, *Cylc1*^−/Y^ *Cylc2*^+/−^ males displayed an “intermediate” phenotype, showing slightly less damaged sperm than *Cylc2*^−/−^ and *Cylc1*^−/Y^ *Cylc2*^−/−^ animals. This further supports our notion, that loss of the less conserved *Cylc1* gene might be at least partially compensated by the remaining *Cylc2* allele.

In general, the evolutionary rate of C-terminal lysine rich region of both Cylicins seems to be highly volatile between conserved and positively selected codon sites, while the lysine residues seem to be strongly conserved. Changes in the C-terminal region, potentially affecting the length of the lysine rich domain, might have an adaptive advantage. Targeted positive selection on codon sites could also be a signature of co-evolution with a rapidly evolving interactor.

Sperm morphological defects and infertility observed in one patient with variants in both Cylicin genes points towards a requirement for human spermiogenesis and fertility. A defect in acrosome formation and the existence of variants in two out of three alleles in both *CYLC* genes is in line with the observations made in mice. The absence of CYLC1 confirmed through immunofluorescent staining indicates a functional impact of the missense variant on the CYLC1 protein. Furthermore, impaired CCIN localization was observed in patient M2270 as well, suggesting that PT has similar roles in human and rodents despite the differences in sperm head shape ^42^. These results suggest that establishing mouse models deficient for Cylicin and other PT proteins might provide insights into mechanisms of human spermiogenesis and cytoskeletal regulation as well. However, with our data we cannot exclude the possibility that there is a discrepancy between mice and men and that the missense variant in *CYLC1* might alone be sufficient to cause the observed phenotype of M2270. The o/e ratios of both genes calculated within the gnomAD database rather indicate a stronger selective pressure on human *CYLC1* (0.08) than on *CYLC2* (0.84). Furthermore, the *CYLC1* variant affects an intolerant region within the protein sequence according to metadome ^43^, whereas the *CYLC2* variant is located in a region that is more tolerant to variation (Figure 7 – supplement 1). Therefore, we cannot definitively validate the hypothesis of an oligogenic disease in men as well.

The *CYLC2* missense variant is inherited by the father of M2270 (Fig. 5A) and, thus, not sufficient to cause infertility. However, the father reported difficulties to conceive naturally. Based on this one family, we cannot exclude an effect of pathogenic heterozygous *CYLC2* variants.

Overall, the identification and detailed characterization of further patients with variants in *CYLC1* and *CYLC2* is warranted to draw firm conclusions on the effect of variants in these genes on spermiogenesis and infertility.

## Acknowledgment

This study was supported by a grant from the German Research Foundation (DFG) to HS (SCHO 503/23-1, project number 458746826), SS (SCHN 1668/1-1, project number: 458746826) and FT (Clinical Research Unit, Male Germ Cells‘, CRU326). We are grateful to Gaby Beine, Angela Egert, Andrea Jäger, Greta Zech, Luisa Meier and Christina Burhöi for excellent technical assistance. We would like to thank the Core Facilities for Microscopy and Analytical Proteomics of the Medical Faculty at the University of Bonn for providing support and instrumentation funded by the Deutsche Forschungsgemeinschaft (DFG, German Research Foundation, project numbers: 388169927, 386936527).

---

**Figure 1 – supplement 1:** Amino acid sequence comparison of CYLC1 and CYLC2 in *C.elegans* and *M.musculus* to *H.sapiens*. KKD/E motifs are highlighted in blue and repeating units are marked by red brackets.

**Figure 1 – supplement 2:** Immunohistochemical staining against CYLC1 and CYLC2 in tissue sections of testis, brain, thymus and spleen. Scale bar: 100 µm.

**Figure 1 – supplement 3:** Immunofluorescence staining against the acrosomal matrix marker protein SP56 (green) and CYLC1 or CYLC2 (red) in round and elongating spermatids. Nuclei were stained with DAPI. Scale bar: 5 µm.

**Figure 1 – supplement 4:** Immunofluorescence staining of CYLC1 and CYLC2 in elongating spermatids of WT, *Cylc1*^−/y^, *Cylc2*^+/−^, *Cylc2*^−/−^, *Cylc1*^−/y^ *Cylc2*^+/−^ and *Cylc1*^−/y^ *Cylc2*^−/−^ mice. Scale bar: 5 μm.

**Figure 1 – supplement 5:** Proteome abundances

**Figure 1 – supplement 6:** Proteome clustering

**Figure 2 – supplement 1:** HE-stained testicular tissue sections of WT, *Cylc1*^−/y^, *Cylc2*^+/−^, *Cylc2*^−/−^, *Cylc1*^−/y^ *Cylc2*^+/−^ and *Cylc1*^−/y^ *Cylc2*^−/−^ mice. Scale bar: 100 µm.

**Figure 2 – supplement 2:** Eosin-Nigrosin staining of epididymal sperm samples from WT, *Cylc1*^−/y^, *Cylc2*^+/−^, *Cylc2*^−/−^, *Cylc1*^−/y^ *Cylc2*^+/−^ and *Cylc1*^−/y^ *Cylc2*^−/−^ mice. Scale bar: 10 μm.

**Figure 2 – supplement 3:** Nuclei of WT, *Cylc1*^−/y^, *Cylc2*^+/−^, *Cylc2*^−/−^, *Cylc1*^−/y^ *Cylc2*^+/−^ and *Cylc1*^−/y^ *Cylc2*^−/−^ sperm stained with DAPI. Scale bar: 5 μm. Elongation and circularity of nuclei from WT, *Cylc1*^−/y^, *Cylc2*^+/−^, *Cylc2*^−/−^, *Cylc1*^−/y^ *Cylc2*^+/−^ and *Cylc1*^−/y^ *Cylc2*^−/−^ sperm. The minimum detection area was set to 1.000 pixels, while the maximum detection area was 7.000 pixels.

**Figure 2 – supplement 4:** Co-staining against CYLC1/CYLC2 (red) and CCIN (green) in epididymal sperm cells of WT mouse. Nuclei were counterstained with DAPI. Scale bar: 2 μm.

**Figure 3 – supplement 1:** TEM micrographs of WT and *Cylc2*^−/−^ sperm and axonemes.

**Figure 3 – supplement 2:** SpermQ analysis of the flagellar beat of WT (green) and *Cylc2*^−/−^ (red) sperm. Average curvature of the flagellum and the arc length are shown.

**Figure 4 – supplement 1:** PNA-lectin immunofluorescence staining of WT, *Cylc1*^−/y^, *Cylc2*^+/−^, *Cylc2*^−/−^, *Cylc1*^−/y^ *Cylc2*^+/−^ and *Cylc1*^−/y^ *Cylc2*^−/−^ testicular tissue. Spermatids at cap phase and acrosome phase of acrosome biogenesis are shown individually. Scale bar: 5 μm.

**Figure 4 – supplement 2:** PAS stained testicular tissue sections of WT, *Cylc1*^−/y^, *Cylc2*^+/−^, *Cylc2*^−/−^, *Cylc1*^−/y^ *Cylc2*^+/−^ and *Cylc1*^−/y^ *Cylc2*^−/−^ mice. Scale bar: 20 µm.

**Figure 4 – supplement 3:** TEM micrographs of degrading damaged spermatids in testicular sections of *Cylc2*^−/−^ mice. Scale bar: 5 µm.

**Figure 5 – supplement 1:** Immunofluorescence staining of α-Tubulin in WT, *Cylc1*^−/y^, *Cylc2*^+/−^, *Cylc2*^−/−^, *Cylc1*^−/y^ *Cylc2*^+/−^ and *Cylc1*^−/y^ *Cylc2*^−/−^ squash testis samples. Spermatids at step 8-9 are shown. Scale bar: 10 μm.

**Figure 7 – supplement 1:**

(A) Localization of the CYLC1 variant found in M2270. The variant affects exon four and an intolerant part of the C-terminal region of CYLC1 according to metadome 41.

(B) CYLC2 variant localization. The missense variant in CYLC2 detected in M2270 affects exon five and a tolerant part of CYLC2 according to metadome 41.

## Methods

### Animals

All animal experiments were conducted according to German law of animal protection and in agreement with the approval of the local institutional animal care committees (Landesamt für Natur, Umwelt und Verbraucherschutz, North Rhine-Westphalia, approval IDs: AZ84-02.04.2013.A429, AZ81-02.04.2018.A369). *Cylc1-* and *Cylc2*-deficient mice were generated by CRISPR/Cas9-mediated gene-editing in zygotes of the hybrid strain B6D2F1. Guide sequences are depicted in Table 2. For *Cylc1*, in vitro transcribed sgRNAs (50 ng/µl each) and Cas9 mRNA (100 ng/µl) (Sigma-Aldrich, Taufkirchen, Germany) were microinjected into the cytoplasm of zygotes as described previously ^44^. *Cylc2*-deficient mice were generated by electroporation of ribonucleoprotein (RNP) complexes into zygotes using a GenePulser II electroporation device (BioRad, Feldkirchen, Germany). For RNP formation, crRNA and tracrRNA (IDT, Leuven, Belgium) were combined in duplex buffer (IDT) to a final concentration of 50 mM each and annealed (95°C, 5 min; cool down to room temperature with - 0,2°C/sec.). For RNP assembly, 4 µM Cas9 protein (IDT) and 2 µM of each annealed cr/tracr RNA were combined in Opti-MEM media (Gibco 11058-021, Thermo Fisher, Carlsbad, USA) and incubated for 10 min at room temperature. RNPs complexes were diluted 1:2 in Opti-MEM, supplemented with 30-40 zygotes in a 0.1 cm gene-pulser cuvette (BioRad) and electroporated (two 30 V square wave pulses, 2 ms pulse length, 100 ms pulse interval). Recovered embryos were cultured over night at 37°C, 5% CO_2_ in G-TL medium (Vitrolife, Göteborg, Sweden) and transferred into the oviduct of pseudopregnant CB6F1 foster mice at 2-cell stage. Offspring were genotyped and gene-edited alleles were separated by backcrossing of founder animals with C57Bl/6J mice. Established mouse lines were registered with Mouse Genome Informatics: Cylc1^em1Hsc^ (MGI: 7341368), Cylc2^em1Hsc^ (MGI:6718280). Mouse lines were maintained as congenic strains on C57Bl/6J background. For all analyses sexually mature males at the age of 8-15 weeks, backcross generation ≥N3 were used.

**Table 2:**
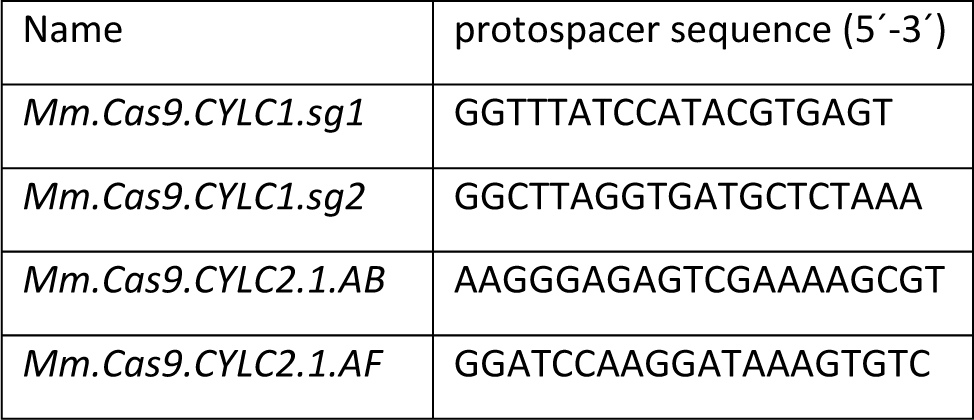
Protospacer sequences.

### Genomic DNA extraction and genotyping

Genomic DNA was extracted from biopsies using the HotShot method ^45^. PCR reactions were assembled according to the manufacturers protocol of the DreamTaq Green DNA Polymerase (Thermo Fisher, EP0712) using gene-specific primers listed in Table 3.

**Table 3:**
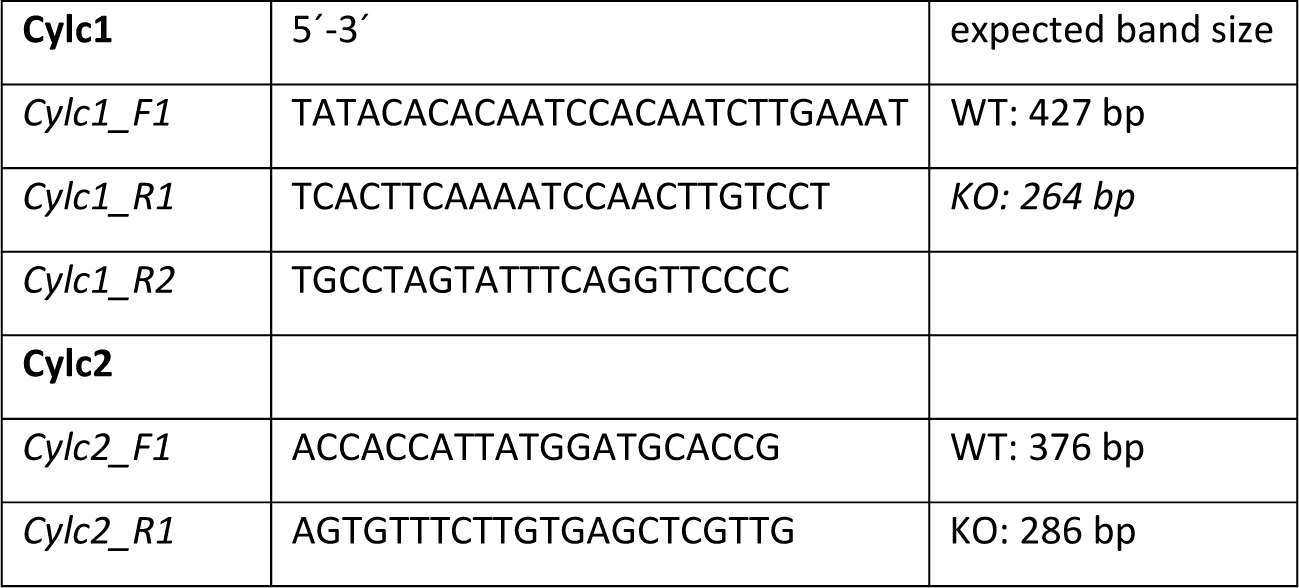
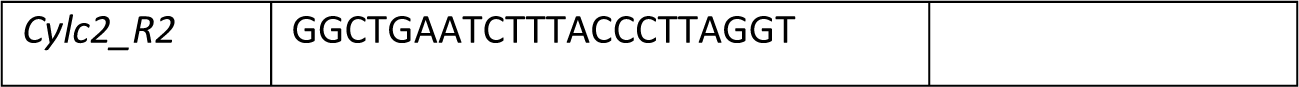
PCR primer sequences.

### Fertility analysis

Males were mated 1:2/1:1 with C57Bl/6J females, which were checked daily for presence of a vaginal plug indicative for successful copulation. Plug positive females were separated and pregnancies as well as litter size were recorded.

### Sampling

Mature sperm were obtained from the cauda epididymis in M2 medium (Merck) or PBS preheated at 37°C. Following several incisions of the cauda, sperm were retrieved by flush-out for 15-30 min. The extracted sperm samples were washed in PBS, centrifuged (4000 rpm, 5 min, 4°C) and snap frozen. For squash tubule samples, fresh testes from three animals of each genotype were transferred to PBS and tubules were dissected as described by Kotaja et al. ^46^. Tubule sections were pressed on a slide, quickly frozen in liquid nitrogen and fixed in 90% ethanol for 5 min.

### Semen analysis

Sperm concentrations for at least 4 animals of each genotype were determined using a Neubauer hemocytometer. Viability of sperm was determined by Eosin-Nigrosin staining as described previously ^47^ for at least 3 animals per genotype. Sperm motility was analyzed referring to the WHO guidelines for analysis of human semen. For all analyses at least 100 sperm per individual were analyzed and the percentage of motile/immotile, viable/inviable sperm was calculated.

### Quantitative reverse transcription-polymerase chain reaction (qRT-PCR)

RNA was extracted from the testis tissue with TRIzol (Life Technologies, Carlsbad, USA; 15596018). RNA concentrations and purity ratios were determined by NanoDrop ONE (Thermo Scientific) measurements. qRT-PCR was performed as described previously ^42^, on ViiA 7 Real Time PCR System (Applied Biosystems) using Maxima SYBR Green qPCR Mastermix (Thermo Fisher; K0221). Replicate of 3 was used for each genotype. Beta-actin was used as a housekeeping gene for normalization. Primers used are shown in table 4.

**Table 4:**
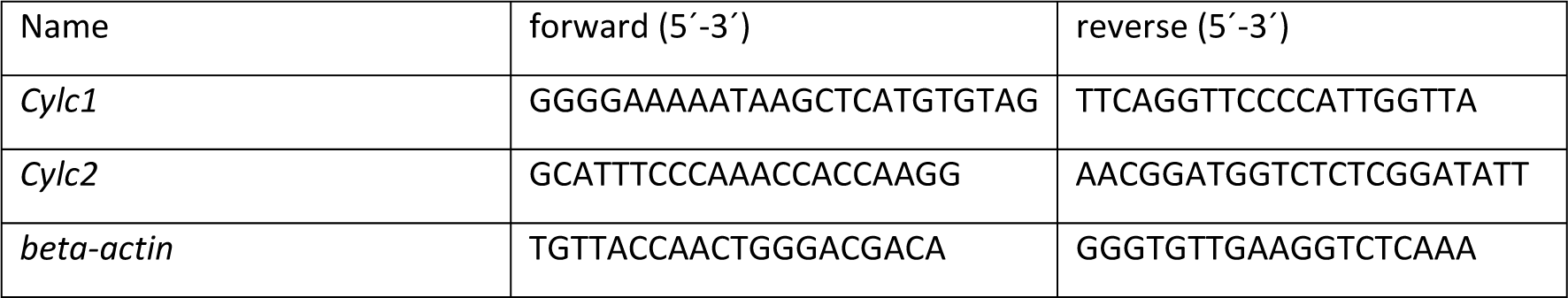
qRT primer sequences.

### Subcellular protein extraction and Western Blot analysis

For the extraction of cytoskeletal protein fraction from sperm cells, Subcellular Fractionation Kit for Cultured cells (Thermo Fisher, #78840) was used according to manufacturer’s instructions, with slight modifications to adjust it to sperm cells. Five subcellular protein fractions were extracted: membrane protein fraction, cytoplasmic protein fraction, soluble nuclear fraction, chromatin bound fraction and cytoskeletal protein fraction. Briefly, epididymal sperm cells were washed in ice cold PBS and centrifuged at 500 x g for 5 min at 4°C. Pellet was resuspended in Cytoplasmatic Extraction Buffer (CEB), incubated for 10 min and centrifuged again at 500 x g for 5 min. Following steps of subcellular fractionation were performed according to manufacturer’s instructions. Quantity of proteins in each fraction was determined using NanoDrop ONE (Thermo Fisher). For further analysis only cytoskeletal protein fraction was used.

Cytoskeletal proteins were separated on 12% SDS gel with 5% stacking gel. Transfer to PVDF membrane was performed using Trans Blot Turbo System (Bio-Rad). Membranes were washed with TBST 1X, stained with Coomassie blue and blocked with 1% milk for 1 h at room temperature with gentle shaking. Primary antibodies anti-CYLC1 and anti-CYLC2 were diluted in milk and incubated overnight at 4°C (for antibody dilutions see table 5). After washing in TBST, membranes were incubated for 1 h at room temperature with polyclonal goat anti-rabbit secondary antibody IgG/HRP (P044801-2; Agilent Technologies/Dako, Santa Clara, 527 CA, United States) diluted 1:2000 in milk. After extensive TBST washing membranes were imaged using WESTARNOVA2.0 chemiluminescent substrate (Cyanagen) or SuperSignal West Femto Maximum Sensitivity Substrate (34095; Thermo Fisher) and ChemiDoc MP Imaging system (Bio-Rad). Membranes were further re-blocked in 3% BSA for 1 h at room temperature with gentle shaking and incubated with α-Tubulin at 4°C overnight. After washing in TBST, membranes were incubated for 1 h at room temperature with polyclonal rabbit anti-mouse secondary antibody IgG/HRP (P0260; Agilent Technologies/Dako) diluted 1:2000 in 3% BSA.

**Table 5:**
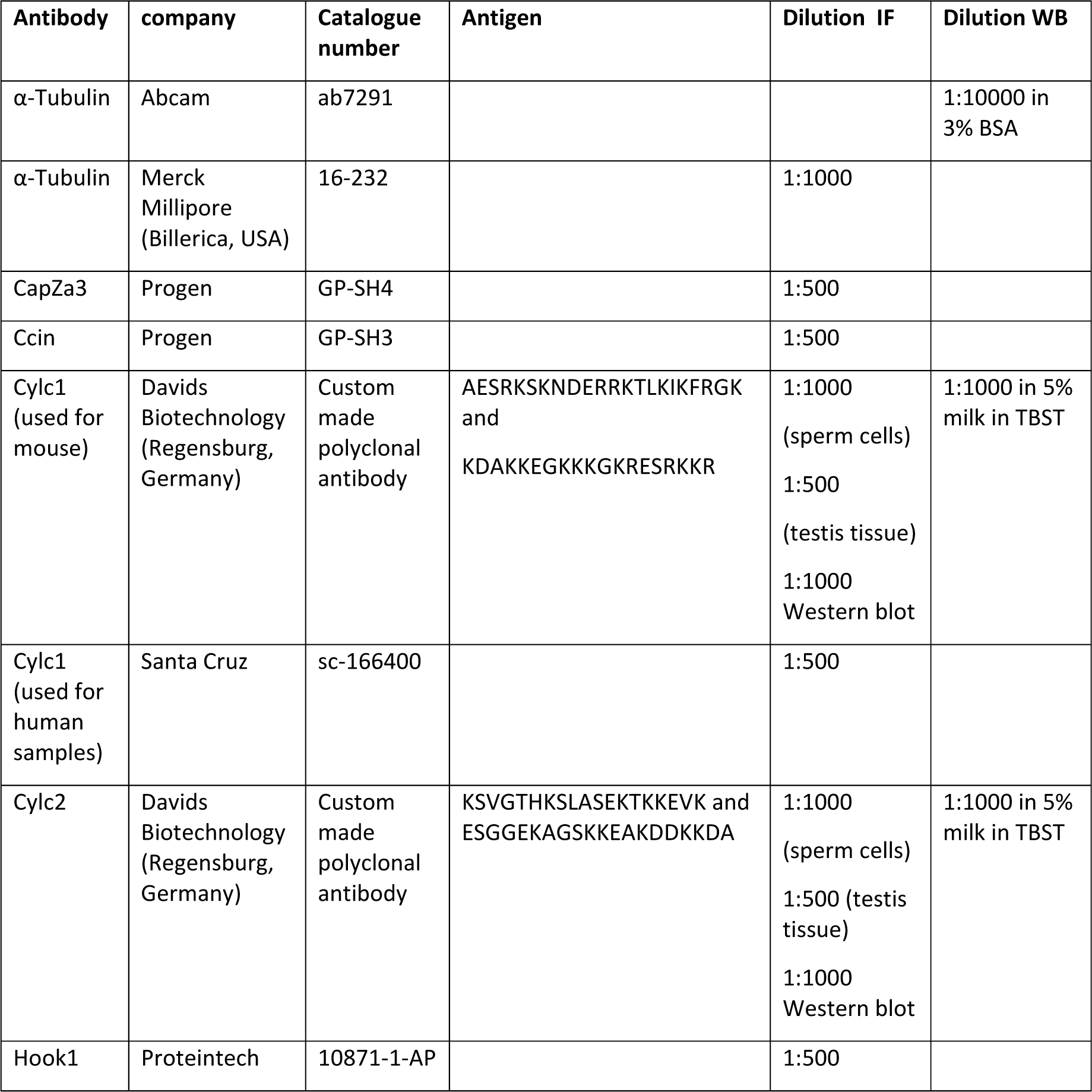
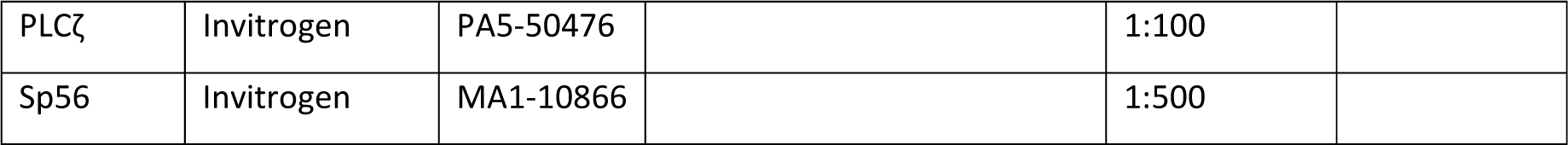
Antibodies.

### Proteomics

#### Peptide preparation

All chemicals from Sigma unless otherwise noted.

Cytoskeletal protein solutions extracted as described in previous paragraph were processed with the SP3-approach ^48^. Briefly, protein lysate with 50 µg protein were subjected to cysteine reduction and alkylation with 20 mM DTT and 40 mM acrylamide in 50 mM triethylammonium bicarbonate (TEAB). Then a mixture of hydrophilic carboxylate-coated magnetic beads (equal amounts of Sera-Mag SpeedBeads, GE Healthcare, cat. no. 45152105050250, and cat. no. 65152105050250) were added at a bead:protein ratio of 10:1 (w/w). Protein binding was induced by adding 3 volumes of ethanol and subsequent mixing for 5 min. Beads with bound proteins were then washed three times with 80% ethanol and finally subjected to overnight tryptic digestion at 37 °C using a trypsin:protein ratio of 1:25. Peptide solutions were separated from the magnetic beads, dried in a vacuum concentrator, and stored at −20°C. Before measurements, 10 µg of peptides were further desalted with C18 ZipTips (Merck Millipore, Darmstadt, Germany) to ensure complete removal of beads.

#### LC-MS analysis

Dried peptides were dissolved in 10 µl 0.1% formic acid (solvent A) Peptide separation was performed on a Dionex Ultimate nano HPLC system (Dionex GmbH, Idstein, Germany) coupled to an Orbitrap Fusion Lumos mass spectrometer (Thermo Fisher Scientific, Bremen, Germany). 1 µg peptides were injected onto a C18 analytical column (400 mm length, 100 µm inner diameter, ReproSil-Pur 120 C18-AQ, 3 µm).

The samples were analyzed by a standard data-dependent (DDA) method: Peptides were separated during a linear gradient from 5% to 35% solvent B (90% acetonitrile, 0.1% FA) at 300 nL/min within 120 min. Data-dependent acquisition was performed on ions between 330 and 1600 m/z scanned in the Orbitrap detector every 2.5 seconds (R=120,000, standard gain control and inject time settings). Polysiloxane (*m/z* 445.12002) was used for internal calibration. z>1 ions were subjected to higher-energy collision induced dissociation (HCD: 1.0 Da quadrupole isolation, threshold intensity 25,000, collision energy 28%) and fragments analyzed in the Orbitrap (R=15,000). Fragmented precursor ions were excluded from repeated analysis for 25 s.

#### Data analysis

Raw data processing of DDA data and analysis of database searches were performed with Proteome Discoverer software 2.5.0.400 (Thermo Fisher Scientific). Peptide identification was done with an in-house Mascot server version 2.8.1 (Matrix Science Ltd, London, UK) against the Uniprot reference proteome for Mus musculus (as of 06/28/23) and a collection of common contaminants ^49^. Precursor ion *m/z* tolerance was 10 ppm, fragment ion tolerance 20 ppm. Tryptic peptides (Trypsin/P) with up to two missed cleavages were searched, propionamide was set as a static modification of cysteines, while oxidation of methionine and acetylation of protein N-termini were set as dynamic modifications. Spectrum confidence of Mascot results was assessed by the Percolator algorithm 3.05 as implemented in Proteome Discoverer ^50^. Spectra without high confident matches (q-value > 0.01) were sent to a second round Mascot search with semi-specific enzyme cleavage and changing the modification of cysteines with propionamide to dynamic. Proteins with two unique proteins in the protein group were reported. For quantification, summed abundances were normalized on total protein amount in Proteome Discoverer.

### High-resolution microscopy of the flagellar beat

Image sequences of mouse sperm were acquired using dark-field at an inverted microscope (IX71; Olympus, Hamburg, Germany), equipped with a dark-field condenser and a high-speed camera (ORCA-Flash4.0 V3, C13220-20 CU, Hamamatsu, Hamamatsu City, Japan). A 10x objective (NA 0.4, UPlanFL; Olympus, Hamburg, Germany) with an additional 1.6x magnifying lens (Olympus, Hamburg, Germany) that was inserted into the light path (final magnification: 16x) was applied. Image sequences were recorded at a rate of 200 frames per second (fps). A custom-made observation chamber was used ^22^. Sperm samples were diluted in THY buffer shortly before insertion of the suspension into the observation chamber. Three WT and three *Cylc2*^−/−^ animals were used.

### Sperm nuclear morphology

For the analysis of sperm nuclear morphology, epididymal sperm samples from 3 animals of each genotype were fixed in methanol and acetic acid (3:1). The samples were spread onto a slide and stained with 4‘,6-diamidino-2-phenylindole (DAPI) containing mounting medium (ROTImount FluorCare DAPI (Carl Roth GmbH, Karlsruhe, Germany; HP20.1)). The sperm cells were imaged at 100x magnification, using a Leica DM5500 B fluorescent microscope. At least 200 pictures were taken from each group and analyzed using Nuclear Morphology software ^51^ according to the developer’s instructions. The minimum detection area was set to 1.000 pixels while the maximum detection area was 7.000 pixels.

### Histology

Bouin fixed testis tissues were paraffinized, embedded and sectioned at 3-5 µm using microtome. For histological analysis, the sections were deparaffinized, hydrated incubated with Periodic acid (0.5%) for 10 min, rinsed with H_2_0 and incubated for 20 min with Schiff reagent. Alternatively, tissue sections were stained with Hemalum solution acid (Henricks and Mayer) and Eosin Y solution (Carl Roth). In both procedures, stained sections were dehydrated in alcohol row and mounted using Entellan (Sigma-Aldrich). Slides were imaged at 20x and 63x magnification under bright field using 5500 B microscope.

### Immunofluorescence/Immunohistochemistry

Bouin fixed testis tissue sections were deparaffinized in Xylene and rehydrated in decreasing alcohol to be used for immunohistochemical staining. Squash testis samples fixed in 90% EtOH were used for the staining of α-Tubulin and HOOK1. For sperm immunofluorescence, mature sperm cells isolated from cauda epididymis were fixed with methanol acetic acid (3:1), dropped on glass slides and air dried. After washing in PBS twice, all samples were permeabilized using 0.1% Triton x100 for 10 min at room temperature. The samples were then blocked with 5% BSA for 30 min, followed blocking with normal horse serum (Vectorlabs, Burlingame, USA; DI-1788) for 30 min at room temperature. For tissue sections, heat-activated antigen retrieval was performed using citrate buffer (pH 6.0). All primary antibodies were incubated over night at 4°C. Antibodies and dilutions are listed in table 5. The respective secondary antibodies were incubated for 1 h at room temperature using VectaFluor Labeling Kit DyLight 488 and DyLight 594 (Vectorlabs, Burlingame, USA; DI-1788, DI-1794). Slides were mounted with DAPI containing mounting medium (ROTImount FluorCare DAPI, Carl Roth; HP20.1).

For IHC staining against CYLC1 and CYLC2, after antigen retrieval and blocking procedures, slides were treated with 6% H_2_0_2_ for 30 min. Slides were then incubated with primary antibodies overnight at 4°C. Biotinylated goat anti-rabbit IgG was used as a secondary antibody and incubated for 1 h at room temperature. Slides were then processed using Vectastain ABC-AP Kit (Vector Laboratories, AK-5001) and stained using ImmPACT® Vector® Red Substrate Kit, Alkaline Phosphatase (Vector Laboratories, SK-5105) according to manufacturer’s instructions. Counterstaining was performed using Hematoxylin.

For the analysis of acrosome biogenesis peanut agglutinin (PNA)-fluorescein isothiocyanite (FITC) Alexa Fluor 488 conjugate (Molecular Probes, Invitrogen, Waltham, USA; L21409) was used on the Buin fixed testis tissues. After permeabilization and blocking, the tissues were incubated with PNA-FITC 5 μg/ml for 30 min at room temperature. Mature sperm were fixed with paraformaldehyde (4%) for 20 min at room temperature. After PBS washing, the sperm samples were incubated with 5 μg/ml PNA-FITC and 5 nM Mito Tracker Red (Cell Signaling; 9082) for 30 min at room temperature. The slides were then mounted with DAPI mount. All stainings were performed on minimum of 3 animals per genotype.

### Transmission electron microscopy

For Transmission Electron Microscopy fresh epididymal sperm and testis tissue were used. After washing with PBS, the samples were fixed in 3% glutaraldehyde at 4°C overnight. The samples were then washed in 0.1 M cacodylate buffer and fixed again in 2% osmium tetroxide at 4% for 2 h. After dehydration in an ascending ethanol row, the samples were contrasted in 70% ethanol 0.5% uranyl acetate for 1.5 h at 4°C. Samples were then washed in propylenoxide, three times for 10 min at room temperature before embedding in Epon C at 70°C for 48 h. Ultra-thin sections were examined using a Verios 460L microscope (FEI) with a STEM III-detector.

### Evolutionary analysis

Evolutionary rates of mammalian Cylicin genes were analyzed according to Lüke et al (2016) ^52^. Briefly, *Cylc1* and *Cylc2* nucleotide sequences were obtained from NCBI genbank and Ensembl genome browser. Phylogenetic trees of considered species were built according to the “Tree of Life web project”. The webPRANK software was applied for codon-based alignment of orthologous gene sequences and results were visualized using the ETE toolkit. To determine evolutionary rates of gene sequences across mammals, for different clades and for individual codon sites, the codeml application implemented in the PAML software was used ^53, 54^. Selective pressures on protein level are represented by calculation of the nonsynonymous/synonymous substitution rate ratio (ω = dN/dS). It distinguishes between purifying selection (ω<1), neutral evolution (ω=1) and positive selection (ω>1) within various models. The M0 model served as basis for all performed analyses. Different codon frequency settings were tested for the M0 model of each gene and the setting with the highest likelihood was chosen. To test whether alternative models describe the selective constraints within a dataset more precise than the M0 model, likelihood ratio-tests (LRT) were performed. Applied models and LRTs are described by Yang ^53, 54^ and Lüke et al. ^52^.

### Study cohort and ethical approval

The MERGE (**M**al**e R**eproductive **G**enomics study) cohort currently comprises over 2,030 men, mainly recruited at the Centre of Reproductive Medicine and Andrology (CeRA) in Münster. The large majority has severe spermatogenic failure, i.e. severe oligozoospermia (<5 Mill./ml sperm concentration), crypto- or azoospermia. So far, only 35 cases were included because of notable sperm morphological defects (≥5 Mill./ml sperm concentration, <4 % normal forms). Common causes for infertility such as oncologic diseases, AZF deletions or chromosomal aberrations were ruled out in advance. Patients with aetiologically still unexplained infertility underwent whole exome sequencing.

All participants gave written informed consent according to the protocols approved by the Ethics Committee of the Medical Faculty Münster (Ref. No. MERGE: 2010-578-f-S) in accordance with the Declaration of Helsinki in 1975.

### Whole exome sequencing and data analysis

After DNA extraction from patients’ peripheral blood lymphocytes WES was performed as previously described ^55^. WES data obtained from 2,066 infertile men was filtered for rare (≤ 0.01 minor allele frequency, gnomAD) variants located within the coding sequence or the adjacent 15 bp of flanking introns in *CYLC1* and *CYLC2*. Patients carrying only one variant in either of the gene were excluded. In case of either bi-allelic *CYLC2* variants or a combination of *CYLC1* and *CYLC2* variants, the whole exome dataset was screened to rule out other potential genetic causes. Variants detected in this study were classified according to the guidelines by the American College of Medical Genetics and Genomics-Association for Molecular Pathology (ACMG-AMP) ^27^ adapted to recent recommendations as outlined in ^56^.

To rule out an alternative genetic cause for the patient’s condition, his exome data was screened for rare (MAF ≤ 0.01, gnomAD and in-house database), homozygous, X-linked or potentially compound heterozygous high-impact variants (stop gain, start lost, stop lost, frameshift, splice site and splice region as well as missense variants with CADD > 20) and rare, heterozygous LoF variants (stop gain, start lost, stop lost, frameshift, splice site) without filtering for a specific set of genes. Respective genes were screened for testis-expression and reports of infertility.

### Sanger sequencing

Sanger sequencing with variant-specific primers was conducted for validation and segregation purposes. The primers to confirm the *CYLC1* (NM_021118.3) variant c.1720G>C p.(Glu574Gln) are 5’-ACTGATGCTGACTCTGAACCG-3’ (forward) and 5’-CCTTCGAGTCACAAAGTTGTCA-3’ (reverse). To confirm the *CYLC2* (NM_001340.5) variant c.551G>A p.(Gly184Asp), the primers 5’-CTGTCGAGGTGGATTCTAAAGC-3’ (forward) and 5’-TGCATCCTTCTTTGCATCCT-3’ (reverse) were used.

### Analysis of the human sperm samples

Human ejaculate samples from healthy donor and patient M2270, were analyzed according to WHO guidelines prior to washing in buffer and centrifuged (1000 rpm, 20 min). The cells were fixed in methanol and acetic acid (3:1) and used for immunofluorescence staining. Samples were dropped on slides and permeabilized with 0.1% Triton x100. After blocking with 5% BSA for 30 min, slides were incubated with primary CCIN or CYLC1 antibodies (concentrations shown in Table 3) for 3 h at room temperature. Secondary antibodies were incubated for 1 h, followed by mounting with DAPI containing medium. All stainings were repeated 3 times using aliquots of the same sample.

### Statistics

For all analyses mean values +/− SD were calculated. Statistical significance was determined by One-way ANNOVA analyses using Bonferroni correction. All experiments were conducted as biological replicates and N is provided in Materials and methods section and/or figure legends.

### Materials and data availability statement

Custom made antibodies used in this study were developed against murine CYLC1 and CYLC2. A limited amount of antibody can be provided upon request.

SpermQ software used for the analysis of the flagellar beat is publicly available https://github.com/hansenjn/SpermQ.

For the purposes of this study a new code was written for color-coded figures and it’s freely available at https://github.com/hansenjn/ColorStackByTimeAndProject

