## supplemental figure for "Cylicins are a structural component of the sperm calyx being indispensable for male fertility in mice and human"

**Figure 1 - supplement 1**

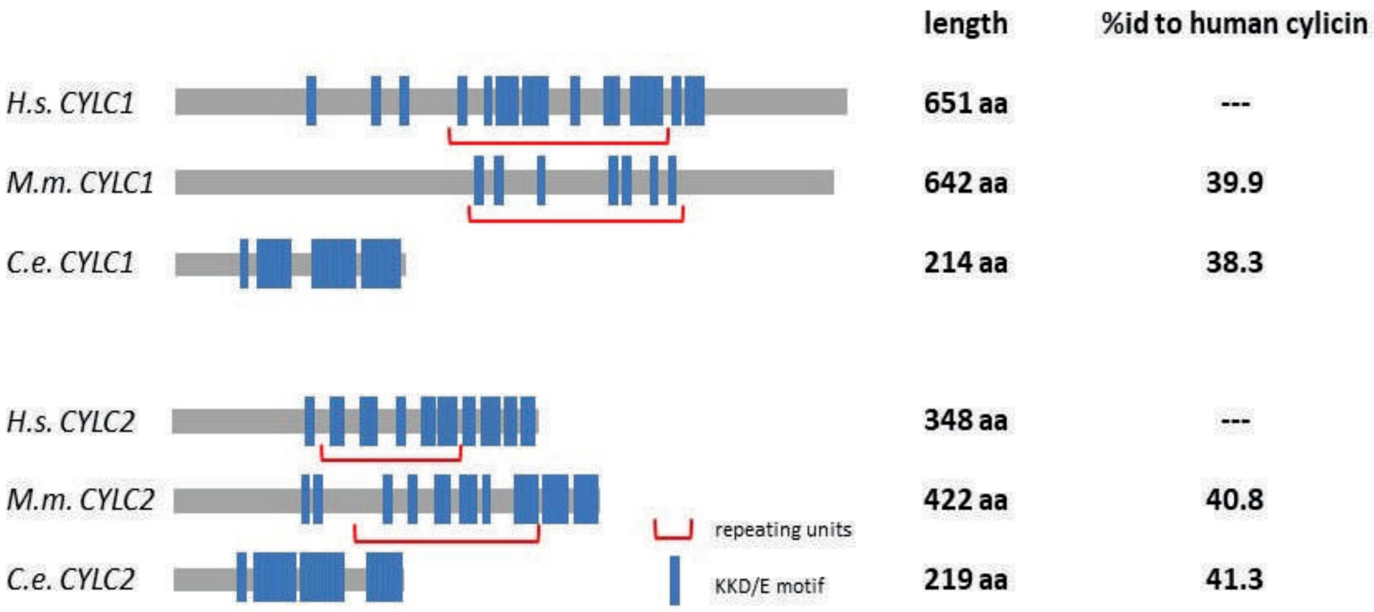

**Figure 1 - supplement 2**

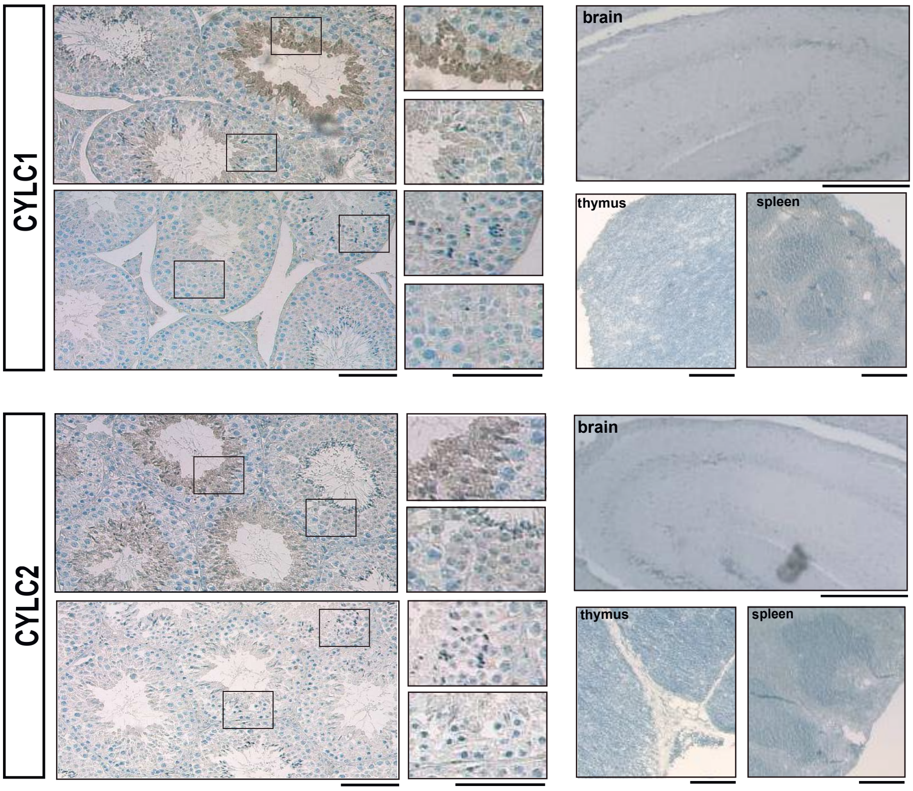

Figure 1 - supplement 3

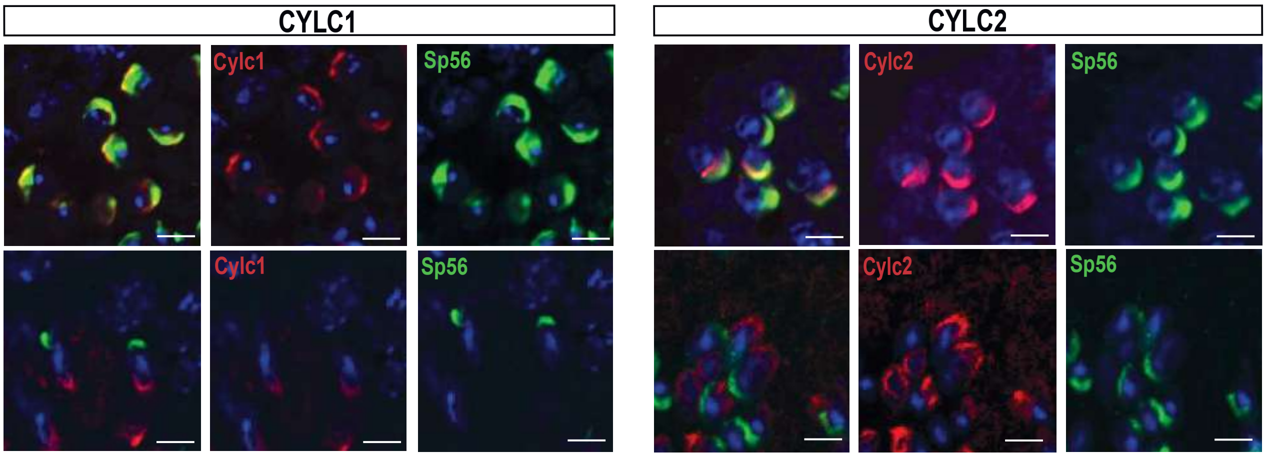

**Figure 1 - supplement 4**

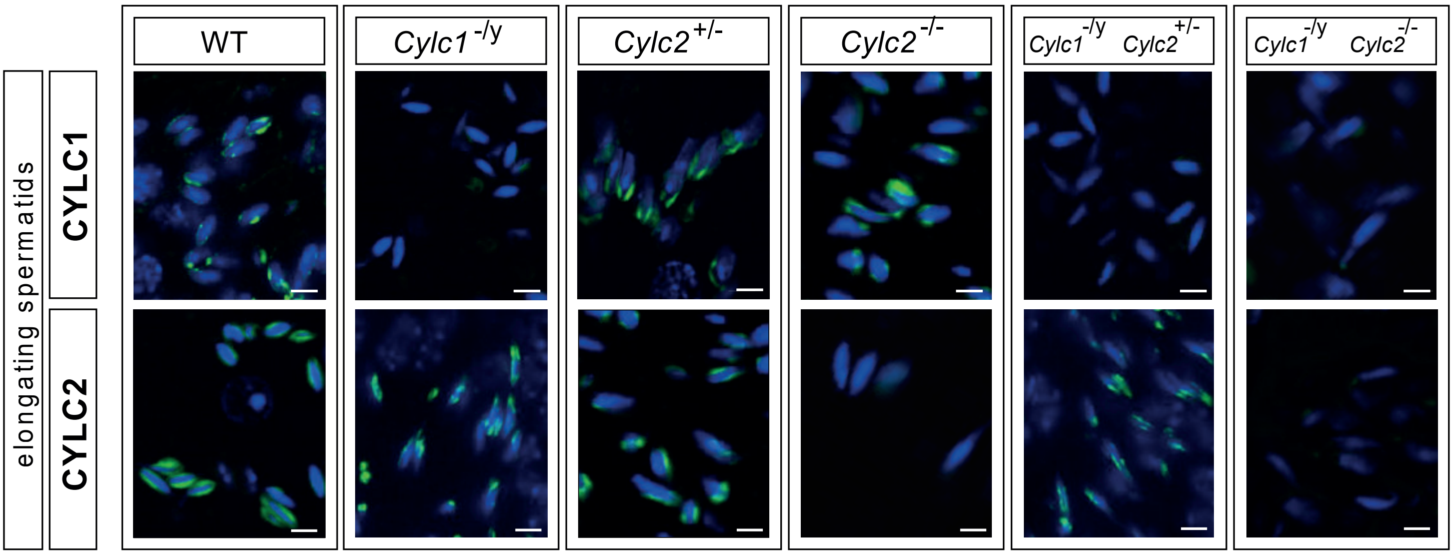

**Figure 1 - supplement 5**

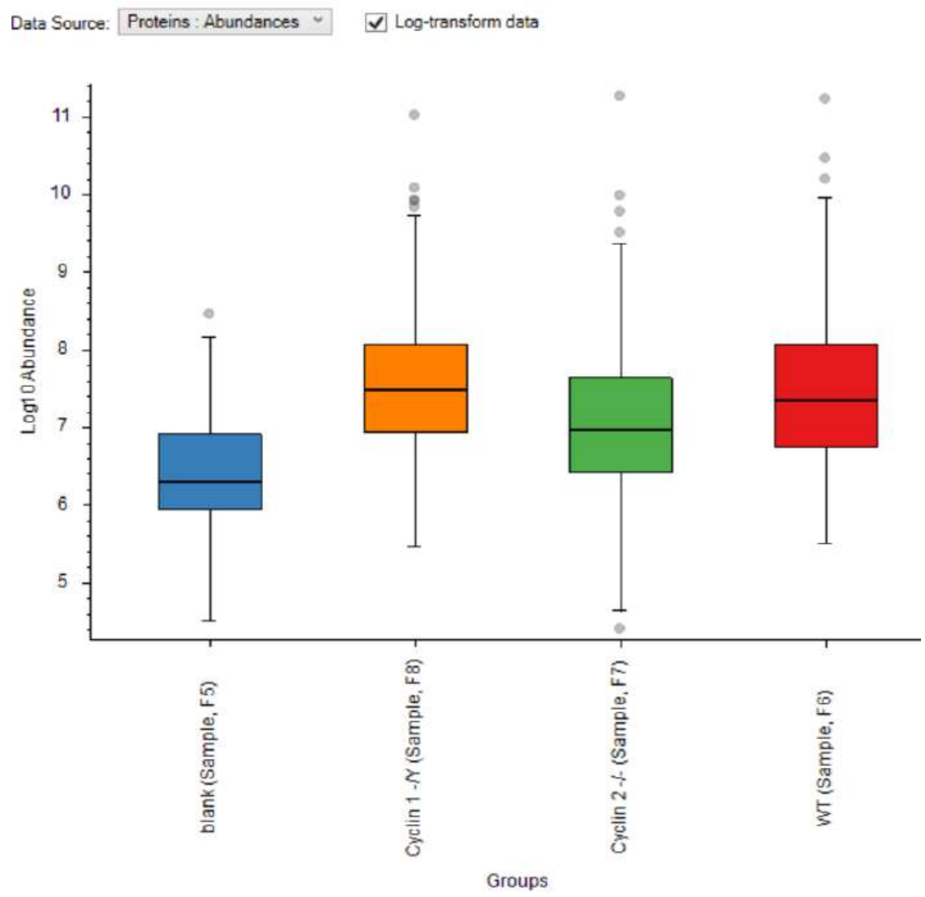

**Figure 1 - supplement 6**

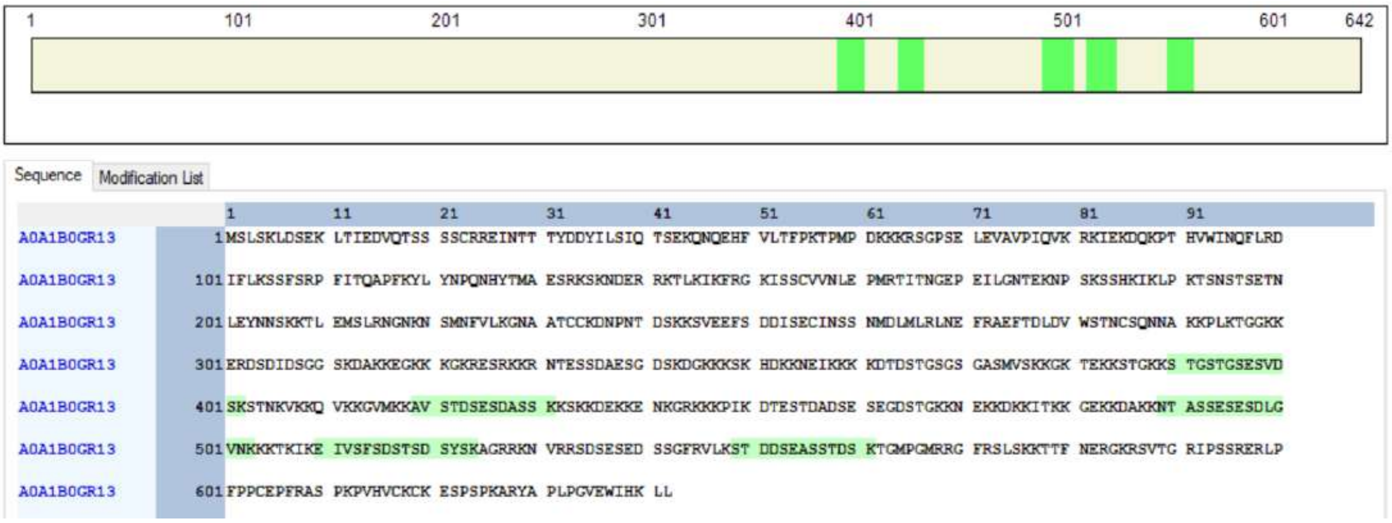


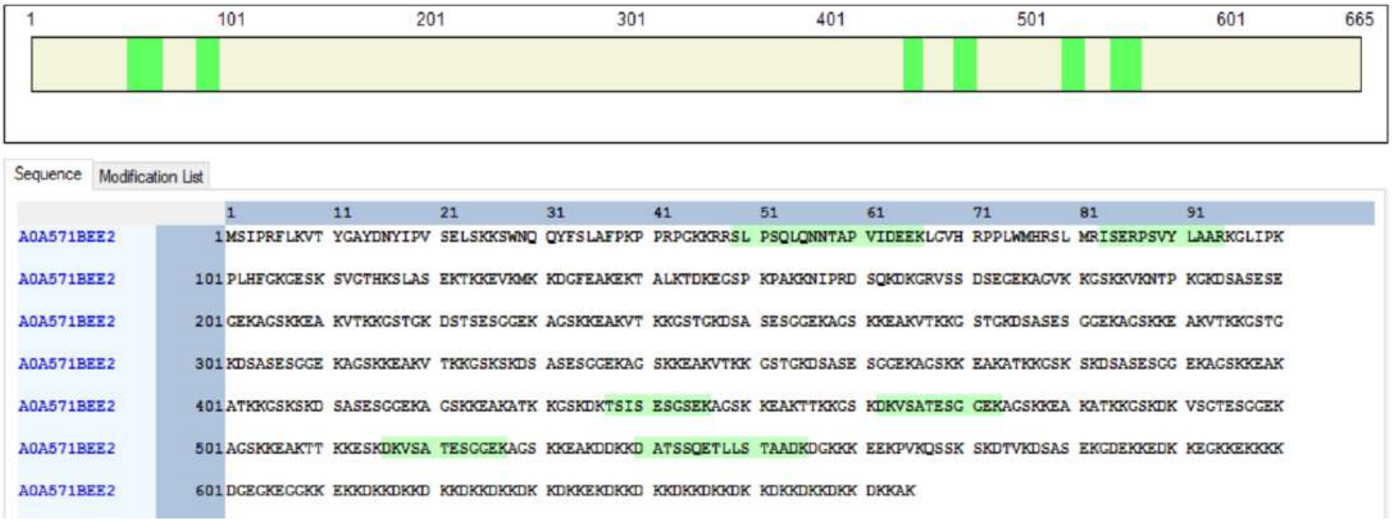

**Figure 2 - supplement 1**

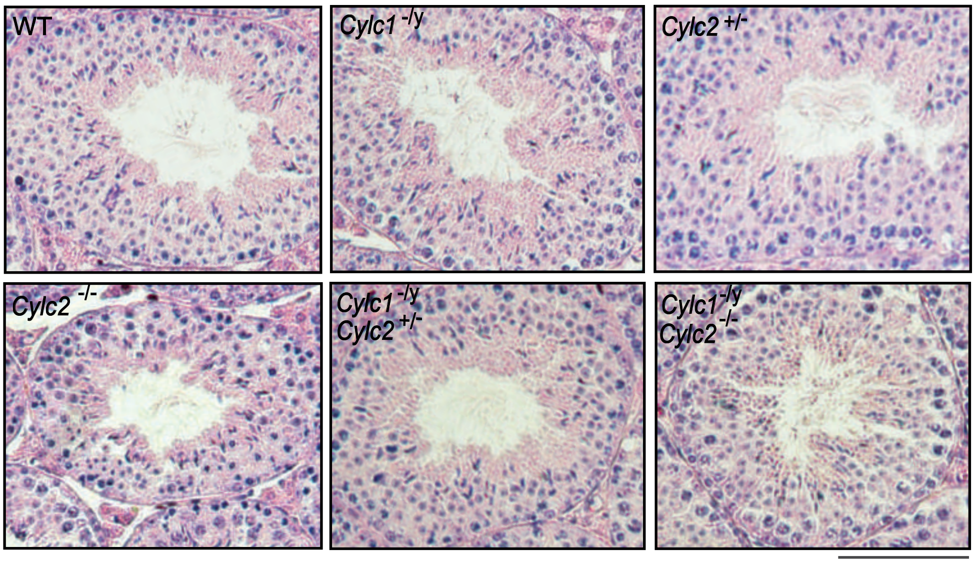

**Figure 2 - supplement 2**

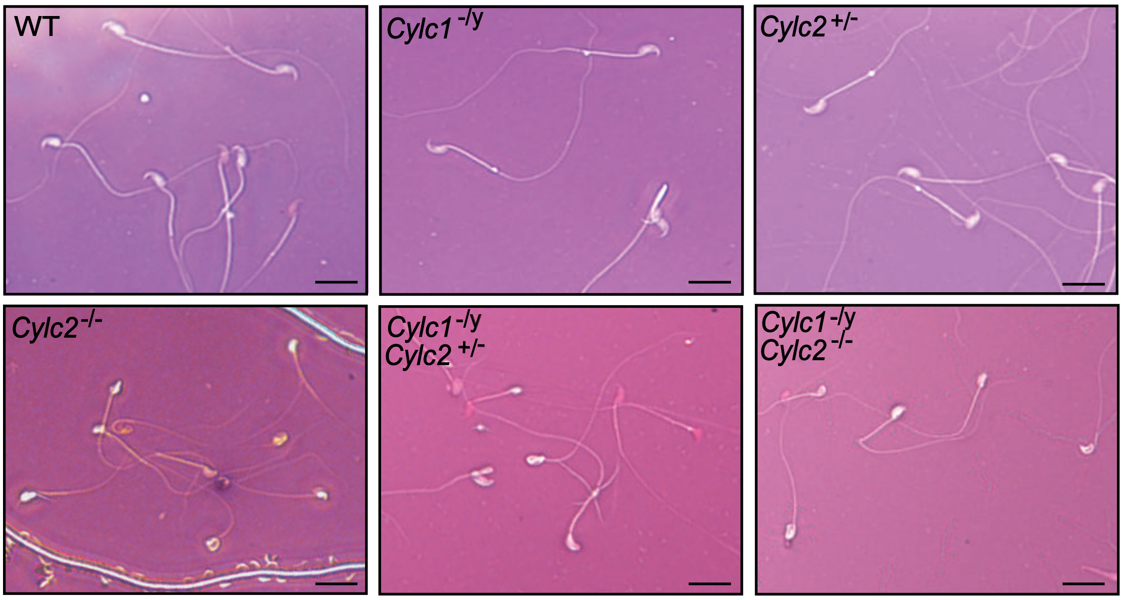

**Figure 2 - supplement 3**

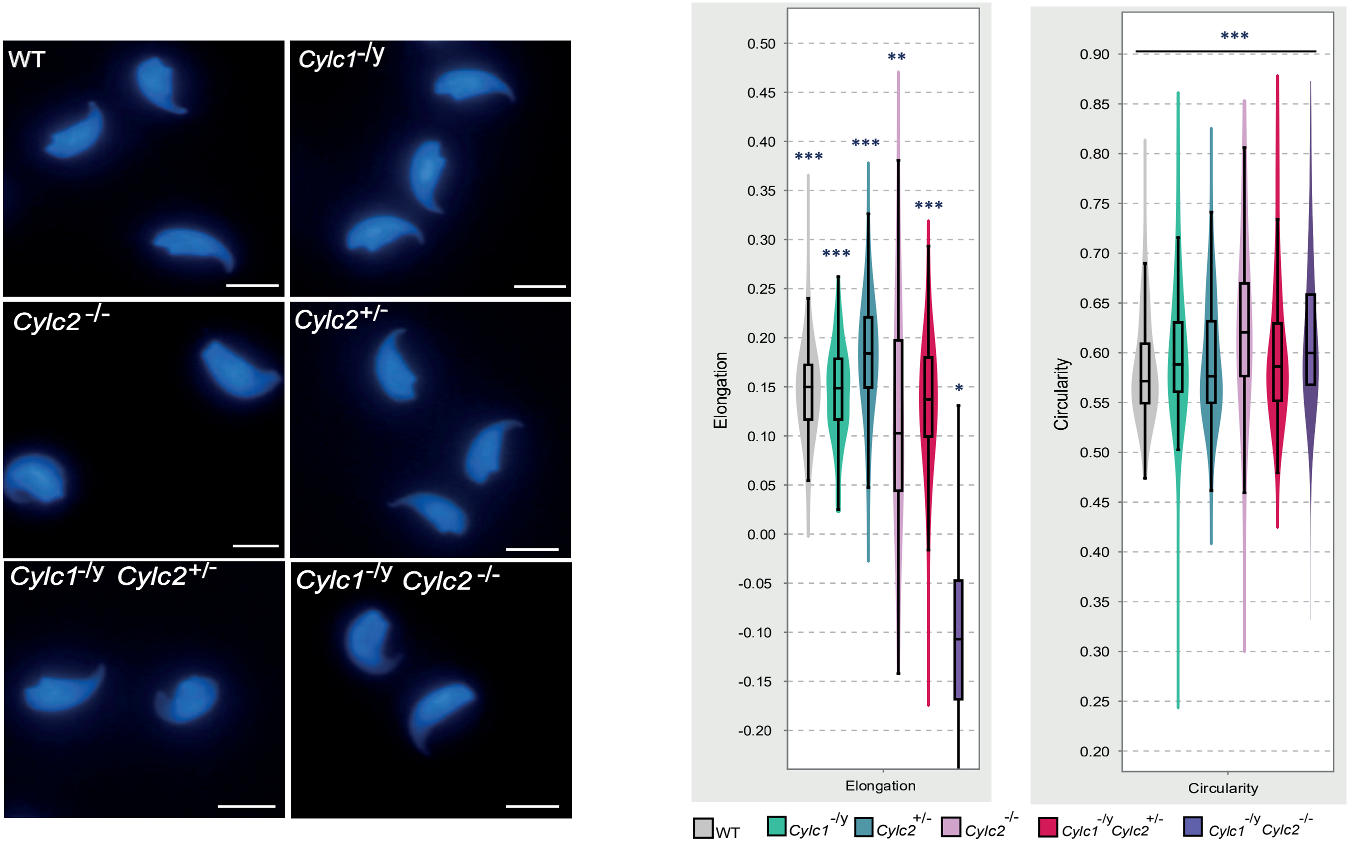

**Figure 2 - supplement 4**

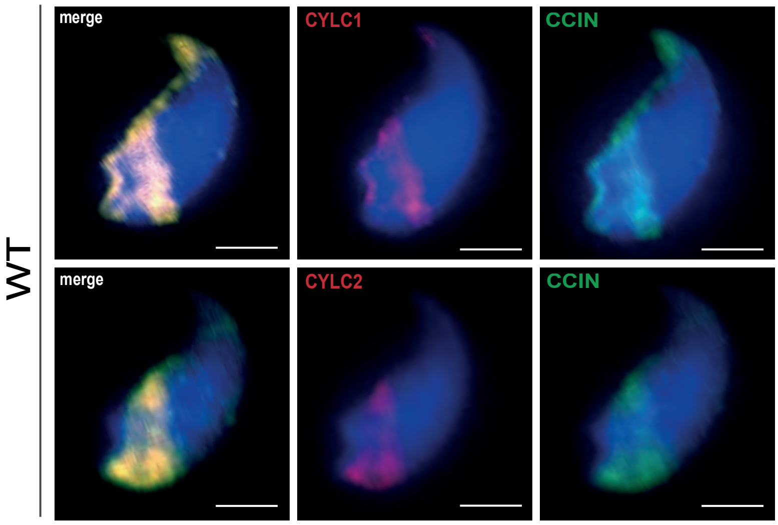

**Figure 3 - supplement 1**

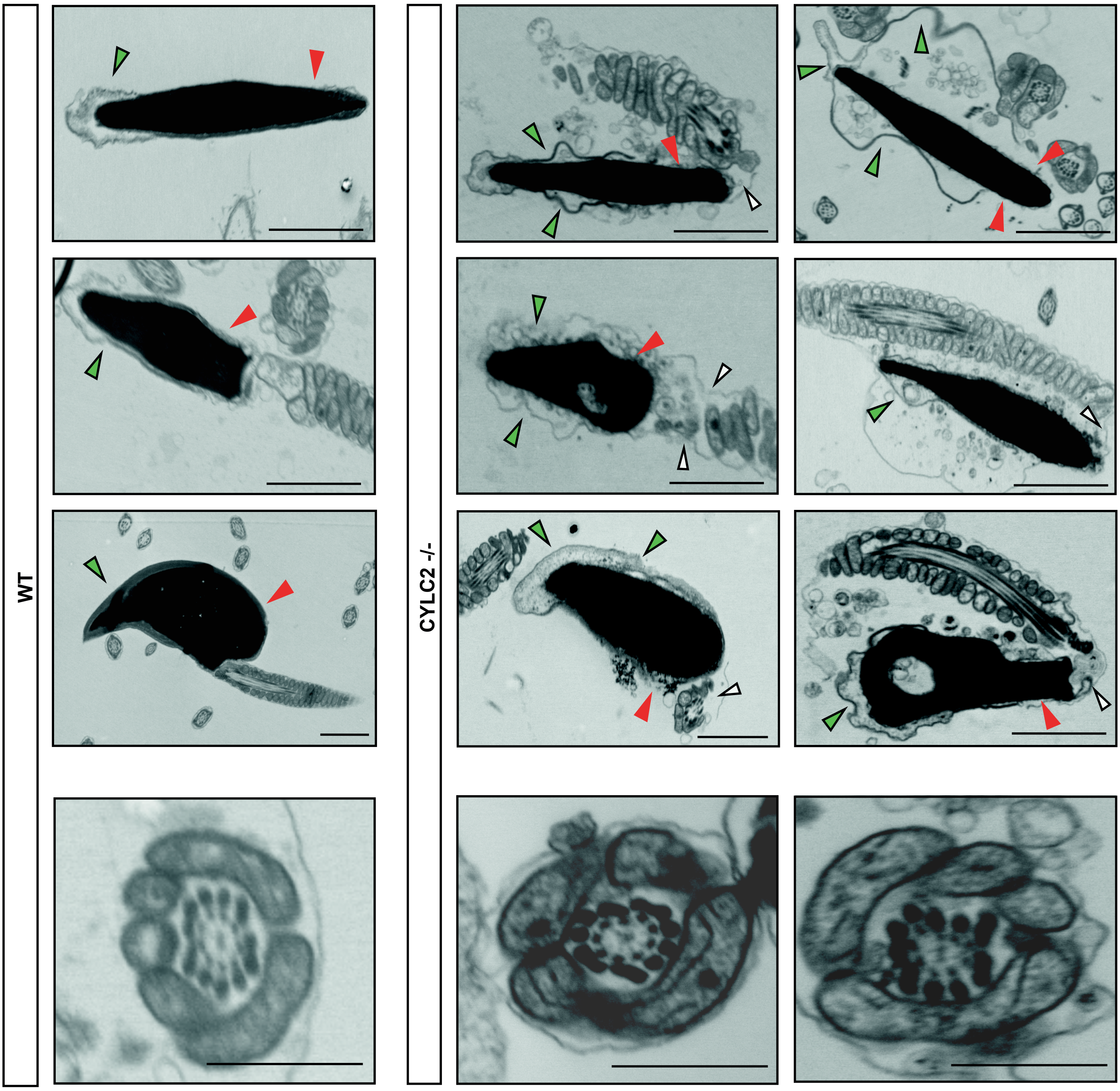

**Figure 3 - supplement 2**

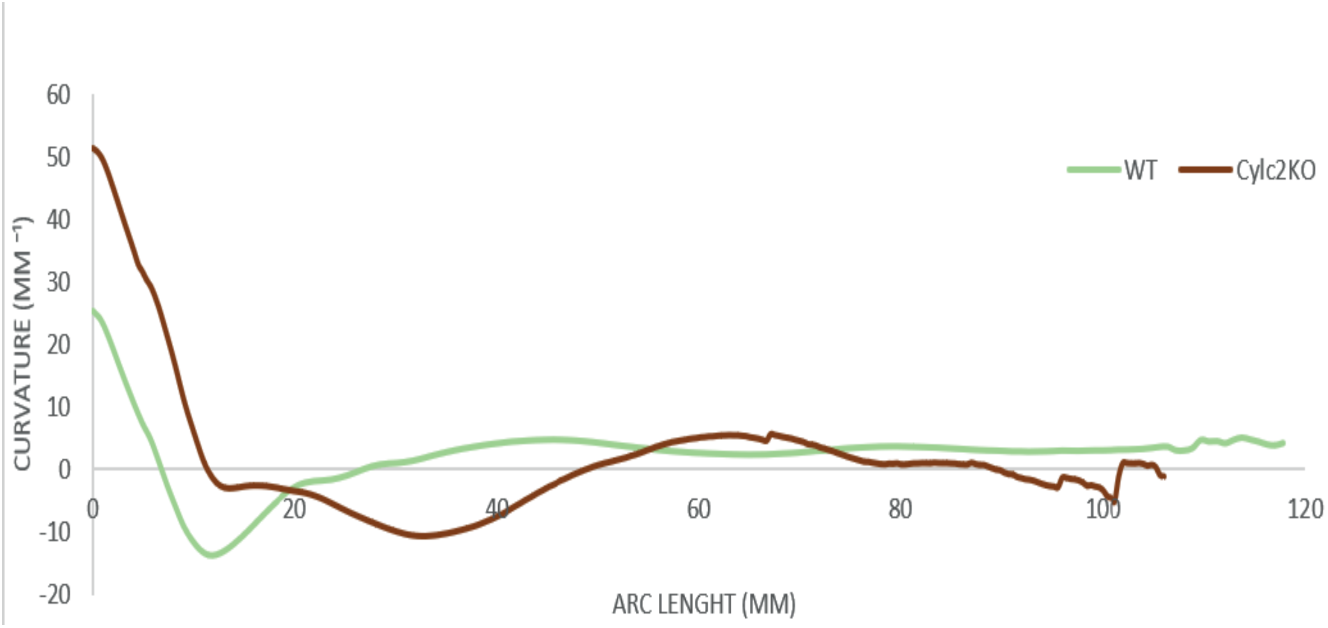

**Figure 4 - supplement 1**

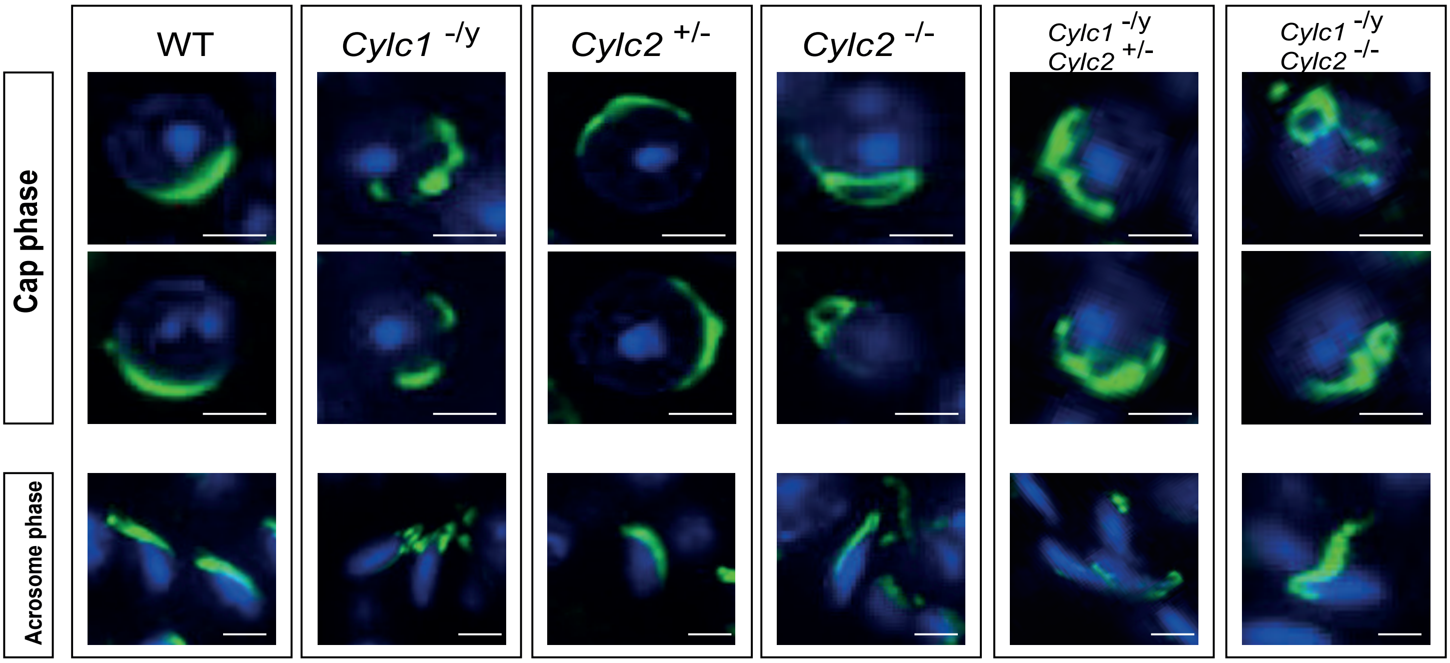

**Figure 4 - supplement 2**

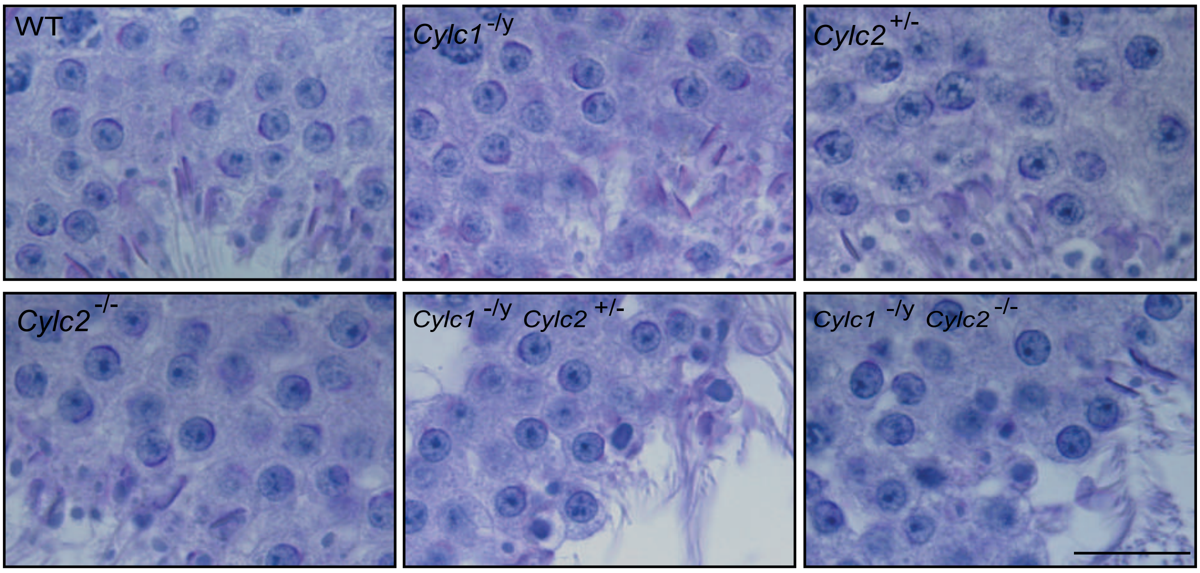

**Figure 4 - supplement 3**

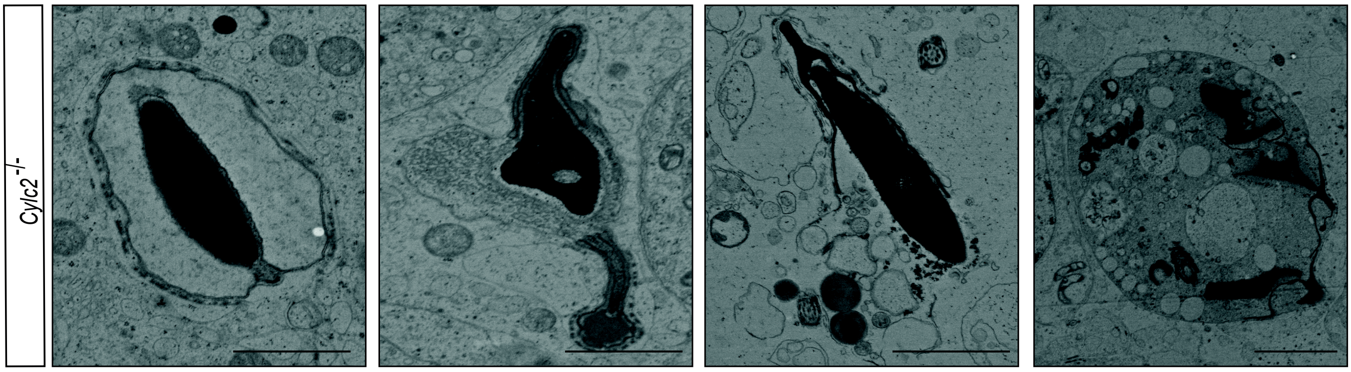

Figure 5 - supplement 1

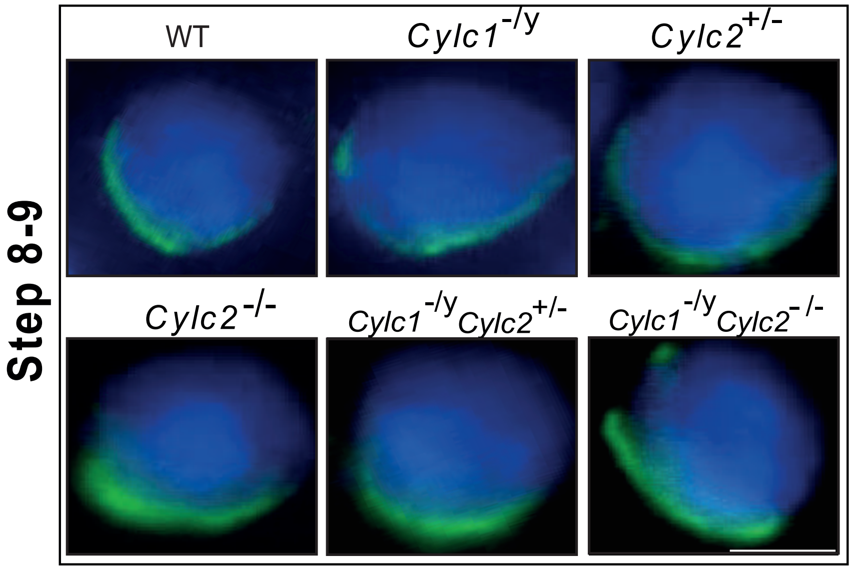

**Figure 7- supplement 1**

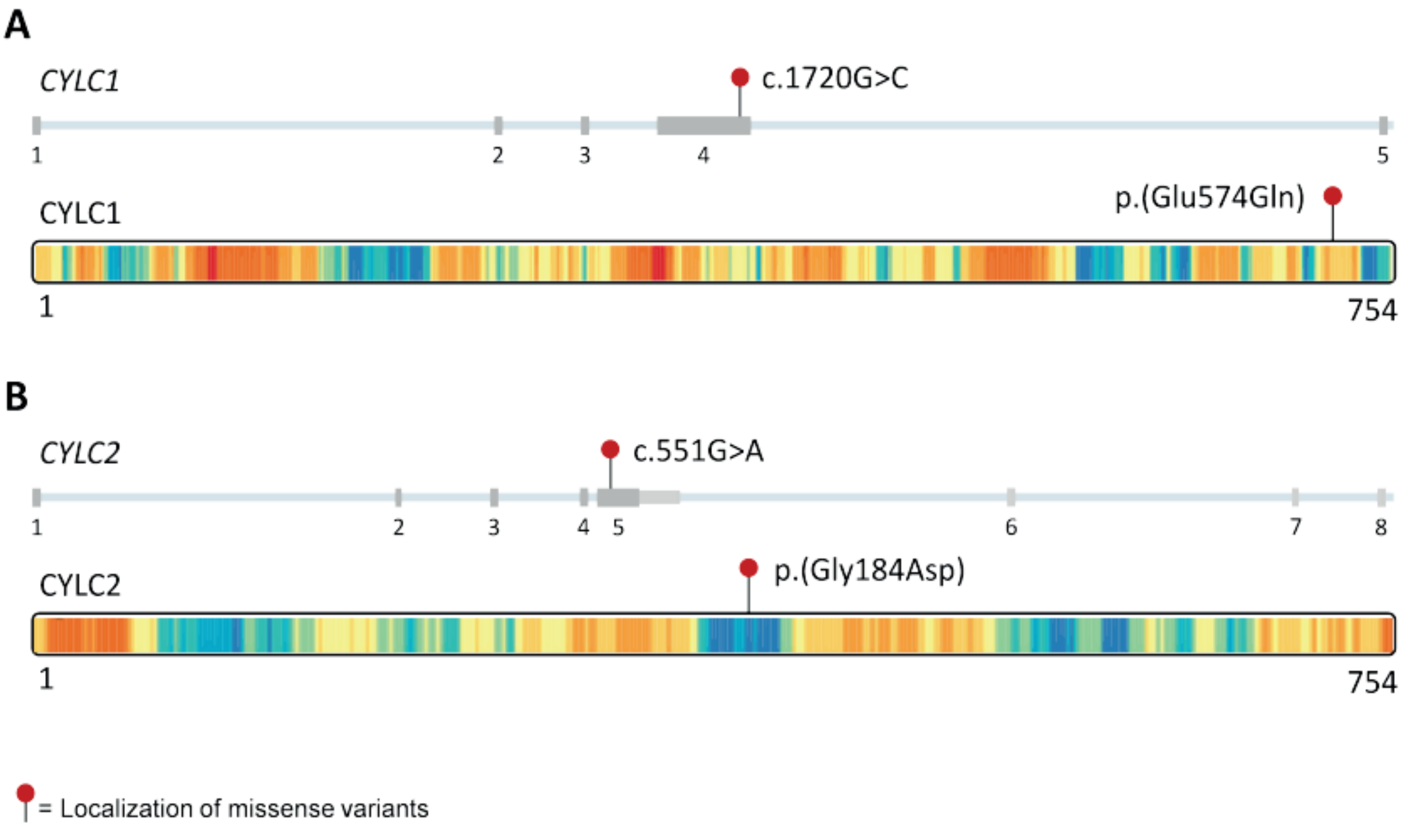
